# UnivAIRRse: A Unified Framework for Organizing and Comparing Adaptive Immune Receptor Repertoire Simulators

**DOI:** 10.64898/2026.02.19.706510

**Authors:** Nika Abdollahi, Sara Kaveh, Sogand Shayesteh, Shayesteh Mommahed, Yasamin Alemzadeh, Reihaneh Zarrin, Fatemeh Chaker Hosseini, Parsa Esmaeili, Reza Hassanzadeh, Sofia Kossida, Changiz Eslahchi

**Affiliations:** Computational Genomics and Immunology (ComputImm) Group, School of Biology, Institute for Research in Fundamental Sciences (IPM), Tehran, Iran; IMGT ®, IGH, Univ Montpellier, CNRS, 34000 Montpellier, France; Department of Engineering Sciences, Faculty of Advanced Technologies, University of Mohaghegh Ardabili, Namin, Iran

**Keywords:** Adaptive immune receptor repertoires (AIRR), Immune simulation, Generative models, Clonal evolution, Benchmarking, Machine learning, Multi-scale modeling, Digital twin frameworks

## Abstract

Adaptive immune receptor repertoire sequencing (AIRR-seq) enables large-scale profiling of B- and T-cell receptor diversity and has become a cornerstone of modern computational immunology. However, AIRR-seq provides only a partial and lossy molecular snapshot of immune dynamics, lacking explicit ground truth for clonal ancestry, lineage trajectories, antigen specificity, and longitudinal immune evolution. This limitation complicates benchmarking, method validation, and mechanistic interpretation of repertoire analysis pipelines.

Here, we introduce UnivAIRRse, a unified hierarchical framework that organizes AIRR simulators within a shared conceptual coordinate system spanning five operational levels, from observed sequence data to the theoretical generative potential of the adaptive immune system. By explicitly distinguishing sequence-, clonal-, specificity-, repertoire-, and generative-level representations, UnivAIRRse enables systematic comparison of simulator assumptions, biological scope, abstraction level, and application focus. To our knowledge, this is the first review to formalize such a unified structure across biological, computational, and functional layers of AIRR simulation.

Using this framework, we review how simulation supports benchmarking, strengthens computational inference, and enables multi-scale investigation of immune repertoire formation and evolution. We identify persistent limitations in existing simulators, including incomplete biological context, limited modularity, restricted interoperability, and overreliance on AIRR-seq as a molecular proxy for complex spatiotemporal immune processes. To operationalize this framework, we provide an interactive web-based AIRR Simulation Landscape Explorer (publicly available at https://www.imgt.org/AIRR-Simulator/) that enables dynamic filtering and comparison of simulators across biological scope, abstraction level, output fidelity, and application focus.

Finally, we outline emerging directions toward digital-twin–ready immune simulation, emphasizing modular architectures, longitudinal multi-omic integration, uncertainty quantification, and dynamic model updating. By providing a coherent conceptual and operational coordinate system, UnivAIRRse establishes a foundation for reproducible, interpretable, and clinically actionable modeling of adaptive immune repertoires, bridging current simulation practices with the next generation of predictive and personalized immunological modeling.

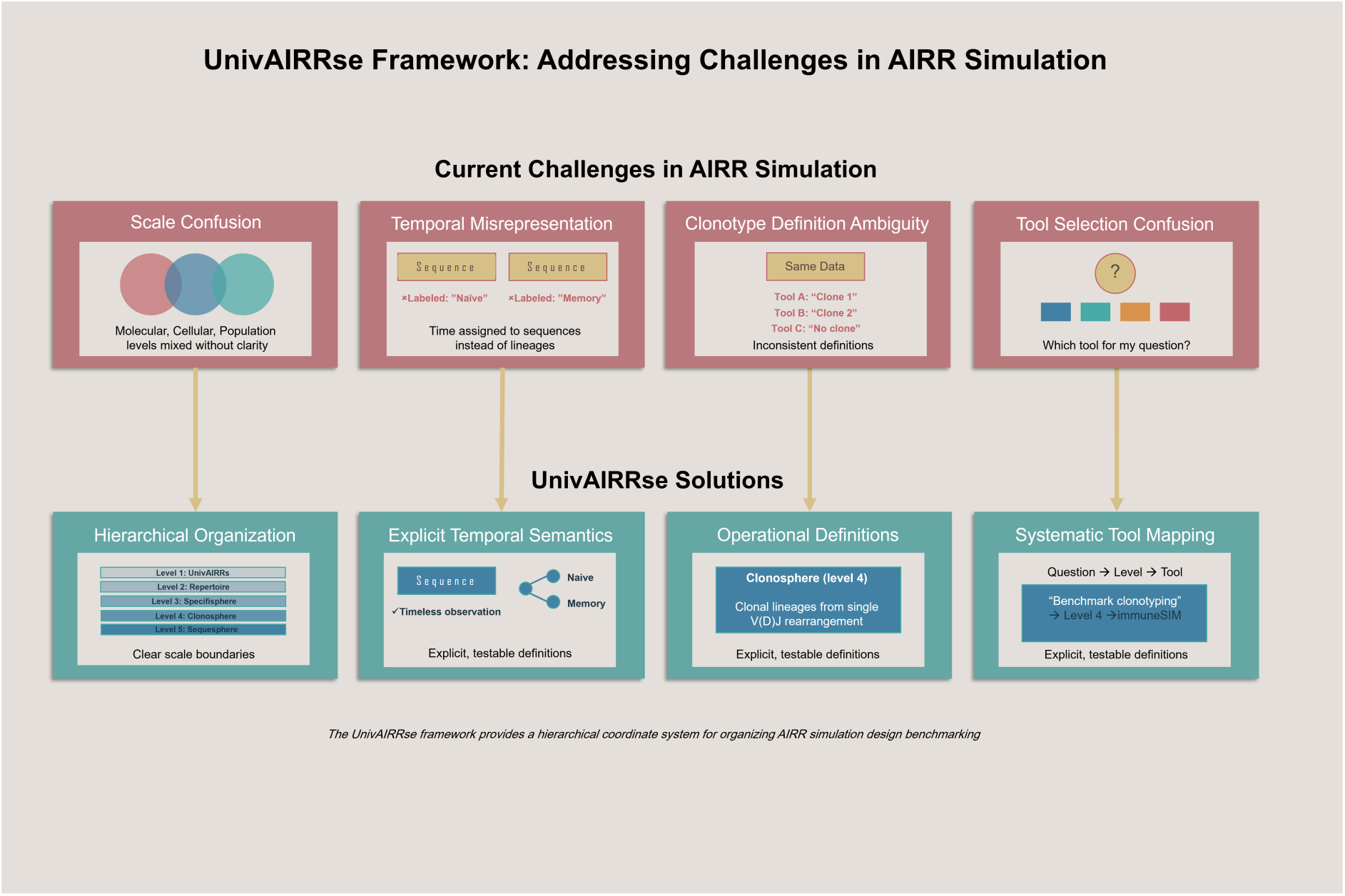

## 1. Introduction

Adaptive immune receptor repertoire sequencing (AIRR-seq) enables high-resolution profiling of B-cell and T-cell receptors and has become a central technique in modern immunology [1, 2]. By generating millions of sequences from individual samples, AIRR-seq provides detailed information on clonal structure [3], somatic hypermutation patterns [4], and immunological history [5]. This depth of measurement has facilitated the study of repertoire diversity, immune responses, and disease-related signatures [6].

Despite these advances, AIRR-seq captures only the observed receptors and not the biological processes that produce them. Experimental datasets lack explicit ground-truth information on clonal ancestry, mutation paths, recombination events, and selection pressures [7, 8]. The absence of this information limits the ability to validate computational methods. Many benchmarking strategies depend on assumptions taken from the same datasets they aim to evaluate, which creates circular reasoning and reduces reproducibility [9, 11] .

AIRR simulators offer a principled solution to this limitation. They model key immunological processes under controlled and interpretable parameters and generate synthetic repertoires with known ground truth [7, 10]. Existing tools simulate V(D)J recombination [7, 8], clonal expansion [2], somatic hypermutation, isotype transitions, and various forms of selection, and they can also include sequencing noise to approximate experimental conditions [12]. These capabilities support hypothesis testing, method development, and performance evaluation in ways that real datasets cannot.

To date, no unified coordinate system has existed for comparing AIRR simulators across biological, computational, and functional layers. The UnivAIRRse framework fills this gap by enabling systematic classification, harmonized interpretation, and benchmark-oriented reasoning across heterogeneous simulation tools.

In this review, we examine the role of simulation within the AIRR research landscape and analyze existing simulators in terms of their biological scope, level of abstraction, and application focus. To provide a unified structure for this analysis, we introduce UnivAIRRse, a hierarchical framework that defines five levels of receptor representation, ranging from concrete sequence data to the full generative potential of the immune system. Using this framework, we trace the evolution of AIRR simulators, highlight their strengths and limitations, and identify challenges that restrict realism, interoperability, and clinical translation. Our aim is to establish a clear and reproducible conceptual foundation for simulation-based research in adaptive immune repertoires.

## 2. Simulation in AIRR Studies: Purpose and Context

Simulation has been central to AIRR research since the earliest developments in computational immunology. Over the past five decades, simulator design has advanced along two major directions. The first trajectory moves toward increasing biological realism, evolving from abstract theoretical models to sequence-resolved and multimodal simulations [13, 15]. The other direction has emphasized computational sophistication, progressing from analytic approximations to probabilistic generative models and later to machine learning frameworks [6, 8]. These advances have shifted simulation from a purely conceptual tool for exploring hypotheses to a source of ground-truth data for benchmarking and, more recently, an integrative platform for modeling immune processes across molecular, cellular, tissue and population levels [15].

### 2.1 Historical Trajectory: From Theory to Data Driven Simulators

Early theoretical models from the 1970s and 1980s, including shape-space formalisms, and immune-network dynamics [13, 9], established core concepts about affinity and selection. However, these models operated at an abstract level and did not produce sequence-level outputs. During the 1990s and early 2000s, bit-string encodings, cellular automata and early agent-based systems such as IMMSIM [16] introduced simplified computational representations of immune population dynamics (see Supplementary Appendix I). These systems were abstract but established the conceptual basis for later mechanistic simulators.

With the introduction of high-throughput AIRR sequencing in the mid-2000s, models began incorporating empirical gene-usage patterns, insertion/deletion distributions, and probabilistic recombination rules [8, 17]. This shift enabled a second generation of simulators around 2016 that integrated AIRR sequence data with explicit recombination and clonal evolution [7, 18]. These tools improved benchmarking and supported more realistic evaluation of inference methods [10].

In the 2020s, development has moved toward multimodal and context-aware systems. Modern simulators integrate machine learning, single-cell transcriptomics, structural information, and spatial data to better capture specificity, cell state and tissue context [6, 19] . This evolution has not been strictly linear. Each generation addressed limitations of earlier models and expanded both biological realism and computational breadth. Meanwhile, AIRR datasets now often include millions of sequences per sample, increasing the need for rigorous ground-truth validation [9]. Even small mismatches between simulator assumptions and downstream analyses can produce misleading conclusions. Simulation has therefore become essential infrastructure for reproducible AIRR research [20].

### 2.2 The Complexity of the Problem: Multi-Scale and Multi-Context Challenges

Effective AIRR simulation must address two interacting challenges. Immune processes unfold across molecular, cellular, tissue, organismal and population levels, and the chosen level of simulation granularity constrains the scientific questions that can be answered [6, 14]. At the same time, researchers often use context dependent simplifications that may be suitable for one study but incompatible with another, complicating comparison and reuse (Mora and Walczak, 2019).

#### 2.2.1 Multi-Scale Biological Complexity

The adaptive immune system operates across several biological levels, each with distinct constraints. Molecular diversity arises from gene usage, sequence composition and biophysical constraints on antigen recognition [6]. Cellular processes involving activation, differentiation and proliferation determine clonal structure and mutation patterns [2, 18]. Tissue-level organization and microenvironmental cues shape compartment-specific repertoires [14]. Organism level processes including immune trafficking, homeostasis, and memory influence longitudinal dynamics [6]. Population-level variation driven by genetics and exposure history affects the distribution of public and private clones [21, 22]. Most simulators represent only part of this landscape, and maintaining consistency across levels remains an open challenge.

#### 2.2.2 Context-Dependent Simplifications

Simulation inevitably requires simplification, but the meaning and implications of these simplifications differ across scientific contexts. Many studies use assumptions tailored to a specific experimental or analytical setting. These assumptions often do not transfer to other settings, leading to inconsistencies and reduced comparability.

A central example is the definition of *specificity*. Depending on the study, specificity may refer to: molecular binding, cellular activation, system level protection, or clinical response. Each definition requires different forms of data for validation [19]. Similar variation appears in *clonotype* definitions, which range from exact sequence identity to inferred ancestry or shared functional behavior. These distinctions can generate markedly different analytical outcomes [4, 18]. *Repertoire* definitions also vary among theoretical, sampled, functional and accessible forms [6, 9], complicating cross-study comparison.

This heterogeneity is also historical. As outlined in Supplementary Appendix I, different generations of AIRR simulators introduced different assumptions and abstractions, many of which remain embedded in contemporary tools. Without explicit clarification, such differences can distort interpretations of diversity, publicness and specificity.

Therefore, simulators must clearly state their assumptions, operational definitions and intended scope. Benchmarking studies must assess whether analytical tools remain robust across alternative definitions and contexts [10, 20]. Such clarity is essential for achieving reliable and reproducible AIRR simulation.

## 3. The UnivAIRRse framework: a hierarchical coordinate system

To organize simulation design and benchmarking, we introduce UnivAIRRse, a hierarchical coordinate system that arranges immune receptor representations into five operational levels [6, 14]. The purpose of this hierarchy is to clarify modeling assumptions, define the biological and technical scope of simulators, and provide a coherent structure for evaluating outputs. As shown in Figure 1, the hierarchy spans a continuum from concrete molecular observations to fully abstract generative potential.

**Figure 1.**
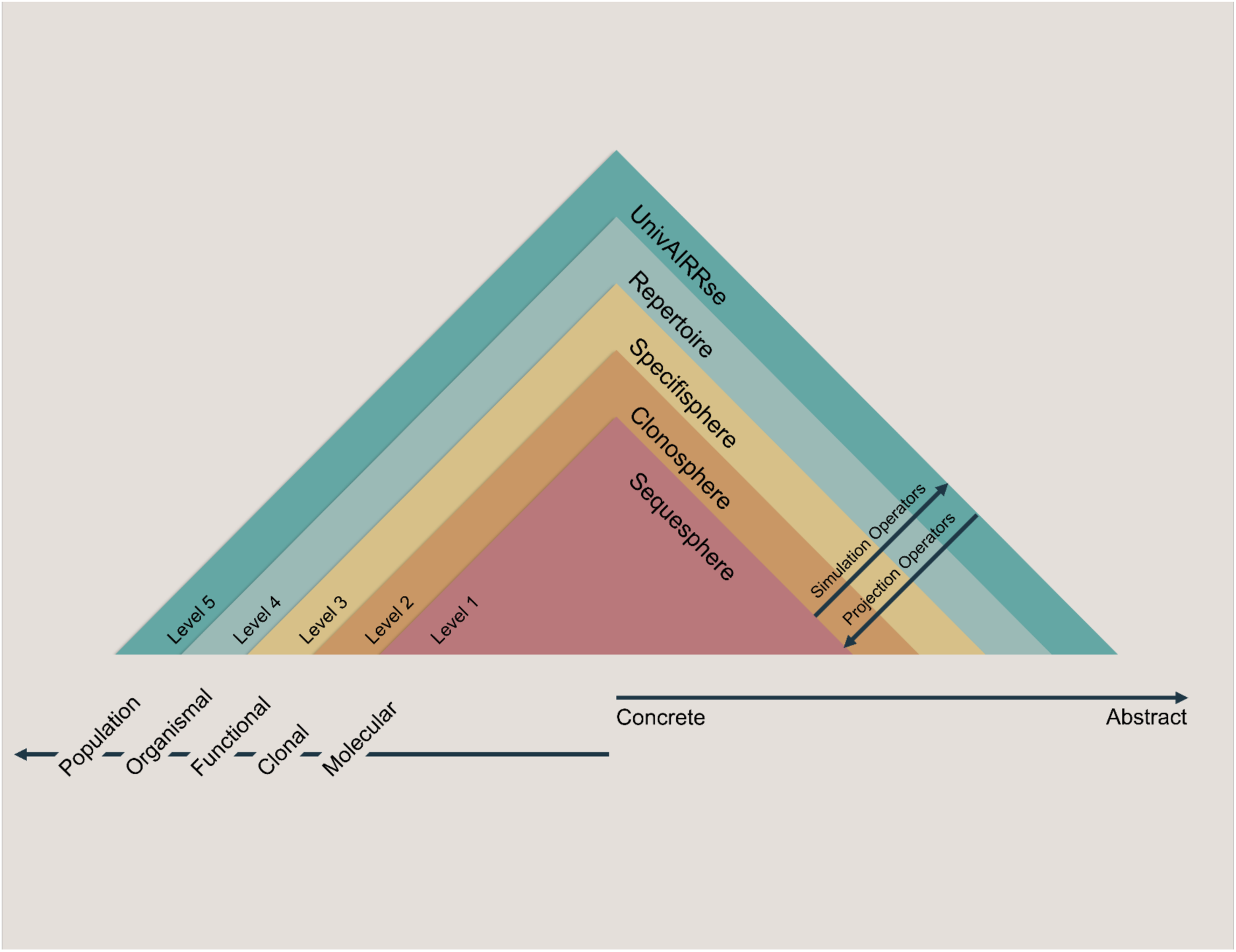
Overview of the UnivAIRRse hierarchical framework. This diagram illustrates the five coordinated levels of immune repertoire representation: the Sequesphere, the Clonosphere, the Specifisphere, the Repertoire, and the UnivAIRRse. The hierarchical ordering (Sequesphere, Clonosphere, Specifisphere, Repertoire, UnivAIRRse) may be conveniently recalled using the mnemonic “S–C–S–R–U”. The levels form a continuum from observable molecular sequence data to increasingly abstract representations of potential diversity. Projection paths show how information moves between levels through summarization, inference, or simulation, providing a unified structure for mapping the scope, assumptions, and outputs of AIRR simulators. Assay and sequencing-error simulators operate in an external measurement layer that acts on sequence-level outputs and therefore are not depicted as part of the five hierarchical UnivAIRRse representational levels.

For ease of reference, the hierarchical progression (Sequesphere → Clonosphere → Specifisphere → Repertoire → UnivAIRRse) can be summarized using the mnemonic “S–C–S–R–U”, reflecting the transition from sequence-level observations to the generative immune universe.

The Sequesphere represents the most concrete level and contains individual sequence records annotated with V and J gene calls, mutation profiles, paired chain information, UMI counts and provenance metadata [7, 10]. The Clonosphere groups sequences derived from a single V(D)J rearrangement and its mutational descendants, capturing lineage structure and clonal evolution [4, 18]. The Specifisphere reflects functional organisation by grouping sequences that recognize a shared antigenic target or epitope, forming specificity neighborhoods [10, 19]. The Repertoire level represents the set of receptors present within an individual or sampled compartment at a specific time point, summarizing diversity, clonal composition, and system level structure [21]. At the highest level, the UnivAIRRse describes the theoretical space of all possible receptors generated by germline gene diversity, junctional processes, and mutational mechanisms [8, 17].

These five levels form a continuous abstraction gradient, moving from directly observable molecular data toward increasingly abstract representations of potential diversity. This structure clarifies the assumptions underlying each simulator and defines the resolution at which outputs should be interpreted. Formal definitions and the projection operators linking adjacent levels are provided in Supplementary Appendix II. Illustrative examples of how these level distinctions affect benchmarking are provided in Box 1.

Figure 1 illustrates this hierarchy. Projection operators summarize or infer higher-level representations from concrete data, while simulation operators generate more complex biological structure. Together, these components form a unified coordinate system for describing simulator scope, identifying biological assumptions and specifying evaluation targets [20].

### 3.1 Time and epistemological gradients in the UnivAIRRse framework

Time is a central challenge in AIRR simulation. Many simulators label sequences as naive, primary, or memory, yet such temporal distinctions cannot be inferred from sequence observations alone [10]. A sequence is a static molecular snapshot captured at a particular sampling moment. Temporal interpretation becomes meaningful only at the clonal level within the Clonosphere, where phylogenetic structure enables identification of naive founders, diversification during primary responses and persistence of memory lineages [2, 18].

At higher organisational levels, such as the Specifisphere and the Repertoire, time appears through population-level changes, including expansion or contraction of antigen-associated neighborhoods and shifts in repertoire composition across longitudinal samples [6]. The UnivAIRRse level is, by definition, time-independent, representing the theoretical generative landscape of possible receptors rather than trajectories realized at specific biological time points.

The framework also moves along an empirical-to-theoretical gradient. The Sequesphere contains directly observable sequence data (nucleotide reads, UMI counts and V/J annotations) shaped by measurement error models [7]. The Clonosphere and Specifisphere represent inferred biological layers and depend on model assumptions, clustering strategies, and threshold choices [4, 6]. The Repertoire level provides integrated descriptors such as diversity indices, gene-usage profiles, and clonal abundances, offering more abstract summaries of immune state. The UnivAIRRse defines the conceptual space of all possible receptors and mutational pathways [8, 17].

Explicitly acknowledging these gradients helps distinguish : what is directly testable, what depends on inference and what belongs to theoretical abstraction. This distinction is essential for interpreting simulator outputs, choosing evaluation metrics, and avoiding conceptual mismatches in benchmarking [9]. The next section applies this framework to classify existing AIRR simulators and highlights the conceptual and technical gaps.

#### Box 1

Practical Applications of the UnivAIRRse Framework for Benchmarking

The UnivAIRRse hierarchy clarifies where ground-truth resides within AIRR data and, in turn, defines how benchmarking should be performed. The examples below show how mismatches between levels can distort evaluation and how awareness of these levels can improve reproducibility.

##### Example 1 | Clonotyping: Sequence Identity vs Shared Ancestry (Level 1 ↔ Level 2)

Clonotyping methods use different definitions. Some group sequences by exact CDR3 and V/J identity (Level 1, Sequesphere), while others infer shared ancestry (Level 2, Clonosphere). Benchmarking must match the intended abstraction level; otherwise, methods may appear to “over merge” or “over split” clones. Paired chain data and dual Level 1-Level 2 labels improve comparability across tools.

##### Example 2 | Antigen Specificity vs Clonal Expansion (Level 3 ↔ Level 2)

Clone size is not a reliable proxy for antigen specificity. Accurate benchmarking of specificity predictors requires true Level 3 labels, such as experimentally validated epitope binders, rather than expansion-based metrics. Expansion alone may reflect proliferation, not functional recognition. Using only level 2 information can mislead model evaluation.

##### Example 3 | Simulator Ground-Truth and Lineage Recovery (Level 1 ↔ Level 2)

Simulators such as immuneSIM, Echidna, AIRRSHIP, and partis produce both: sequence-level labels (Level 1), and true clonal lineages (Level 2). This dual annotation supports rigorous evaluation of clonotyping and phylogenetic reconstruction using: precision/recall, tree distance metrics, or measures of Clonosphere accuracy.

##### Example 4 | Toy Repertoires: Matched Diversity, Divergent Clonality (Level 4 ↔ Level 2)

Two repertoires may show identical diversity indices (Level 4) but having very different clonal structures (Level 2). Combining Level 2 lineage metrics with Level 4 clone size distributions

### 4. Existing Simulators: Landscape and Categorization

AIRR simulators differ widely in their biological assumptions, modeling granularity, and intended purposes. The UnivAIRRse framework provides a structured way to organize this heterogeneity. It compares simulators along three key dimensions: 1) the biological processes they model, 2) the abstraction level at which they operate, and 3) the application focus guiding their design. Together, these dimensions offer a coherent map of simulation capabilities and clarify the assumptions embedded within each framework.

#### 4.1 Categorization by Biological Process

Simulators can first be grouped by the biological processes they reproduce. V(D)J recombination oriented simulators such as repgenHMM [8], IGoR [7] and OLGA [16] operate in the Sequesphere.

They generate rearranged sequences and provide descriptors including generation probabilities and event-level logs of recombination [7, 8]. These tools support estimation of baseline generation probabilities, constructing synthetic naive repertoires, and providing priors for downstream models.

Clonal expansion and somatic hypermutation frameworks include immuneSIM [10], AIRRSHIP [20], AbSim [2], Echidna [23] and partis [18]. They simulate clonal identifiers, B cell lineage trees, and somatic hypermutation trajectories [4, 18]. These frameworks enable benchmarking of clonotyping, lineage reconstruction and phylogenetic inference.

Selection-oriented simulators such as SONIA [24], soNNia [24], Ruiz Ortega [21] and Böttcher [1] model post-recombination selection pressures. They predict public and private clone distributions within or across individuals and support comparison of repertoires at the cohort scale [21].

Specificity-oriented tools such as TAPIR [19] and TULIP [25] model receptor–antigen interactions. They provide functional binding scores and support translational tasks including antigen prediction and target discovery [19].

Evaluation-oriented frameworks, including sumrep [9], compute summary statistics, divergence metrics and diagnostic profiles used to assess similarity between real and simulated repertoires [9].

Measurement-focused simulators such as InSilicoSeq 2.0 [12] introduce platform-specific sequencing noise and assay bias [12]. They support stress testing of analytical pipelines and evaluation of error models.

A complete mapping of these process-based categories to their corresponding UnivAIRRse domains and use cases is provided in Table 1, which groups representative AIRR simulators according to the biological mechanisms they model, the domains they occupy within the UnivAIRRse framework and the types of ground-truth labels they generate. This Table serves as a concise map of the simulation landscape, highlighting how different tools align with recombination processes, clonal and SHM dynamics, selection and publicness modeling, antigen-specificity prediction, statistical evaluation and assay or read-error simulation.

**Table 1.** Categorization of AIRR simulators by biological process. This Table groups representative AIRR simulators by the primary biological processes they model and maps them to their dominant UnivAIRRse domain. Stable IDs (in parentheses) allow cross-referencing with Appendices III and IV (Tables S2–S10). “Typical outputs / ground truth” summarizes the labels or annotations available for benchmarking. “Example use cases” illustrates how each simulator class supports methodological evaluation and mechanistic investigation.

| Category | Representative Tools (Stable ID) | Primary UnivAIRRse Domain(s) | Typical Outputs / Ground Truth | Example Use Cases |
| --- | --- | --- | --- | --- |
| **VDJ recombination simulators** | IGoR [7] , OLGA [16] , repgenHMM [8] | Sequesphere | Generated rearrangements, generation probabilities, recombination event logs (partial truth) | Estimating generation probabilities, creating synthetic naive repertoires, providing priors for selection models |
| **Clonal expansion and SHM frameworks** | immuneSIM [10], AIRRSHIP [20] , AbSim [2] , partis paired , SHM models [4] , Echidna [23] | Clonal, Repertoire | Clonal identifiers, simulated mutation histories and clone-size dynamics, phylogenetic tree structures (not explicit biological lineage trees), and per-cell clone labels for single-cell data | Benchmarking clonotyping and phylogenetic reconstruction, modeling affinity maturation dynamics, and evaluating multi-omics, and single-cell pipelines |
| **Selection and public clone models** | SONIA , soNNia [24], Ruiz Ortega model [21] , Böttcher model [1] | Specifisphere, Population | Post selection probabilities, public clone metrics, cohort level sharing predictions | Quantifying selection, modeling public and private clones, comparing repertoires across cohorts |
| **Antigen specificity models** | TAPIR [19] , TULIP [25] | Specifisphere | Predicted receptor epitope assignments, functional binding scores (no sequence level ground truth) | Target discovery, epitope generalization, computational triage prior to experimental validation |
| **Evaluation and statistical frameworks** | sumrep [9] | Repertoire | Summary statistic profiles, divergence metrics | Comparing real and simulated repertoires, reproducibility assessment, quality control of analytical pipelines |
| **Assay and read error simulators** | InSilicoSeq 2.0 [12] | Primary UnivAIRRse Domain(s): *External measurement / assay layer (acts on Sequesphere outputs; not part of the hierarchical levels in Fig. 1)* | Platform specific error profiles, amplicon biases, coverage distributions | Stress testing error models, evaluating UMI or PCR bias, benchmarking end to end sequencing workflows |

With the exception of AbSim, most simulators listed in this category do not generate explicit biological lineage trees reflecting true germinal-center ancestry. Even AbSim simulates phylogenetic trees rather than mechanistic lineage trees. Therefore, these frameworks primarily support benchmarking of phylogenetic reconstruction rather than full biological lineage reconstruction.

#### 4.2 Categorization by Abstraction Level

Abstraction level provides a second perspective for comparing AIRR simulators. Each class of tools focuses on a distinct layer of immune system organisation and contributes specific types of ground-truth or modeling structure.

Molecular-scale simulators such as repgenHMM and OLGA focus on nucleotide rearrangement. They reconstruct recombination outcomes and report generation probabilities together with event-level annotations.

Clonal-scale frameworks such as AIRRSHIP and Echidna capture lineage structure, clonal identifiers and patterns of somatic mutation reflecting affinity-maturation dynamics.

Within the repertoire scale, tools such as immuneSIM and SONIA summarize aggregate repertoire-level behaviour through clone-size distributions, diversity metrics, and selection-related features.

At the functional scale, tools such as TAPIR and TULIP estimate antigen or epitope specificity by linking receptor sequence or structure to functional interaction.

At the population scale, models by Ruiz Ortega and Böttcher [21] quantify clone sharing, publicness metrics and cross-individual variation.

The final layer concerns measurement processes. Read generators such as InSilicoSeq 2.0 [12] replicate platform-specific noise and biases to create synthetic data with known error profiles. A consolidated mapping of these abstraction layers to their UnivAIRRse domains, expected outputs and major use cases appears in Table S2 (Appendix III), which summarises this abstraction based classification by aligning AIRR simulators with the corresponding modeling layers. This overview clarifies how each simulator operationalizes biological mechanisms at a specific computational scale and highlights the assumptions, outputs and intended use cases associated with each abstraction level. Relationships between these layers, benchmarking requirements and structural limitations are detailed in Supplementary Appendix III.

#### 4.3 Categorization by Application Focus

Application focus provides a third perspective for comparing AIRR simulators. Each class of tools is designed around a specific analytical objective, shaping assumptions, outputs and abstraction level.

Benchmarking-focused simulators (immuneSIM, Echidna) generate defined ground truth, enabling evaluation of: gene assignment, clonotyping, lineage reconstruction, single-cell integration, and noise sensitivity. A detailed summary of benchmarking metrics and performance comparisons appears in Supplementary Table S3 (Appendix II).

Personalisation-oriented frameworks (repgenHMM, SONIA) reconstruct individual-specific priors: selection coefficients, probabilistic generation landscapes, or personalized repertoires. Representative tools for each application class appear in Supplementary Table S2 (Appendix II).

Prediction and translational simulators (TAPIR, Ruiz-Ortega) extend simulation toward clinical or functional interpretation, aiding antigen prediction, public and private clone analysis and identification of disease associated repertoire signatures. Exploratory and educational tools (immuneSIM, partis) provide intuitive environments for visualizing clonal evolution, affinity maturation and repertoire wide structure. These tools are often used for hypothesis generation, teaching and conceptual demonstration.

Across these applications, simulators share several structural limitations. Recurrent issues include incomplete contextual modeling, nontunable biological assumptions, limited availability of ground truth, restricted provenance tracking and weak integration with multiomics or machine learning workflows. Table S5 (Appendix II) links application focus with these limitations, offering a unified summary.

Having introduced the three major dimensions that structure the AIRR simulator landscape,, we now turn to how the UnivAIRRse framework can be used in practice. Rather than adding a new taxonomy, the following components operationalize these dimensions by highlighting integration-oriented simulators, providing a concise decision guide (Box 2) for choosing an appropriate simulator, and outlining common hallmarks of successful frameworks (Box 3). Together, these elements translate the conceptual structure established in Section 3 into practical guidance for selecting, designing, and advancing AIRR simulation tools.

##### Box 2

How to Choose a Simulator: A Short Decision Guide

With the growing diversity of AIRR simulators, choosing the most appropriate framework has become a nontrivial but critical task. The following four-step guide supports transparent and reproducible selection:

1. **Define target level(s):** sequence, clonal, specificity, or population scope.
2. **Match capabilities to metrics:** e.g., lineage trees for clonal validation, P_gen_ for generative likelihoods, or receptor–epitope predictions for translational studies.
3. **Assess sampling and assay models:** account for biases, sequencing depth, and error profiles.
4. **Plan validation:** integrate simulated and empirical datasets, evaluating higher-order repertoire features such as lineage topology, diversity indices, and clonal overlap

#### 4.4 AIRR Simulation Landscape: An Interactive Web Explorer

To provide a practical interface for navigating the AIRR simulator landscape, we developed AIRR Simulation Landscape, an interactive web-based explorer publicly accessible via the IMGT® portal (https://www.imgt.org/AIRR-Simulator/).

The interface renders a two-dimensional map in which the vertical axis corresponds to species and the horizontal axis corresponds to publication year. Users can interactively filter simulators through a category switcher that includes approach, biological scope, chain support, output fidelity, use-case focus, and time-series capability. Each filter activates a color-coded legend, and hovering over any simulator reveals a standardized tooltip summarizing its modeling scope, typical outputs, chain support, interface type, species coverage, and intended applications.

This interactive system serves as an operational extension of the UnivAIRRse framework by enabling rapid identification and comparison of simulation tools, for example, locating HMM-based recombination models, simulators with native time-series support, or paired-chain models for lineage inference. A representative view of the interface is shown in Figure 3. Technical implementation details are provided in Supplementary Appendix V.

**Figure 2.**
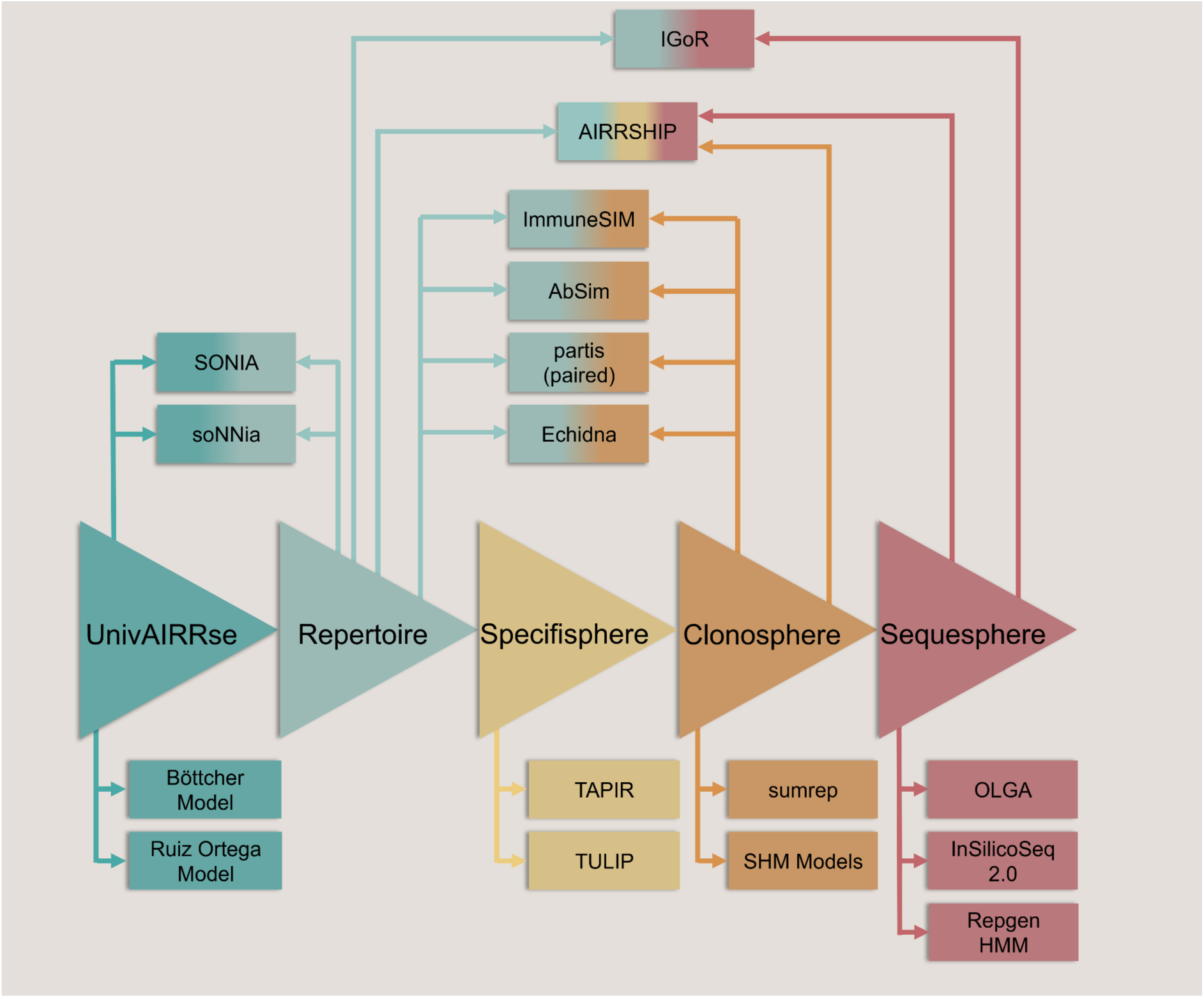
Positioning of AIRR simulators within the UnivAIRRse framework. Mapping of representative simulators onto the hierarchy, following the same right-to-left ordering used in Figure 1 (Sequesphere → Clonosphere → Specifisphere → Repertoire → UnivAIRRse). Tools are positioned according to their dominant biological scope (from molecular recombination to population-level dynamics), their abstraction level, and their application intent. Overlaps reflect the multi-scale and multi-purpose nature of modern AIRR simulation frameworks.

**Figure 3.**
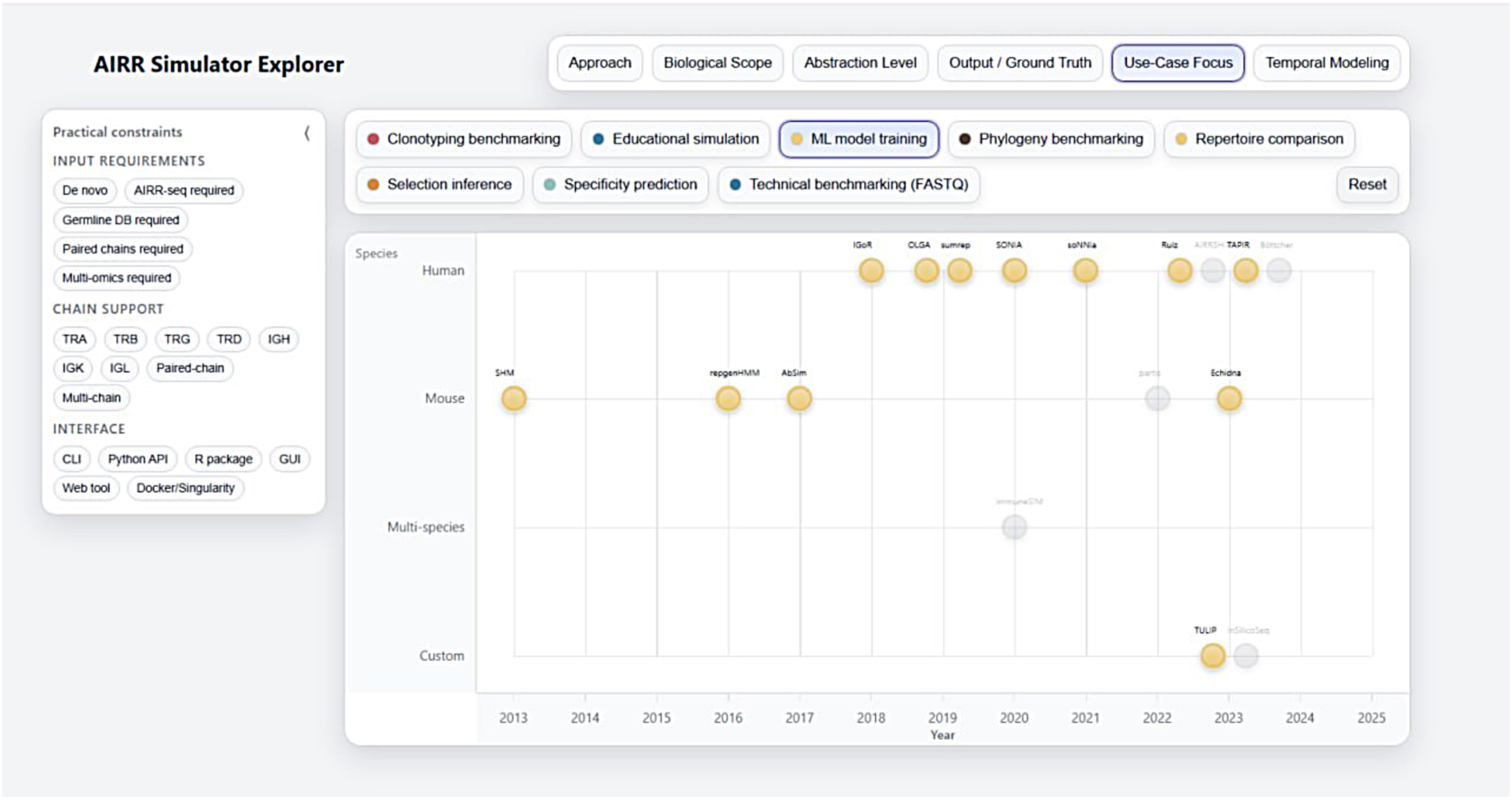
AIRR Simulation Landscape Explorer. Representative view of the interactive web-based AIRR Simulation Landscape Explorer, publicly accessible via the IMGT® portal. Each simulator is positioned on a two-dimensional map according to species (y-axis) and publication year (x-axis). The interface allows dynamic filtering across multiple dimensions, including modeling approach, biological scope, abstraction level, output/ground-truth fidelity, use-case focus, and temporal modeling capability. Color-coded legends highlight selected categories, while interactive tooltips provide standardized summaries of each simulator’s modeling assumptions, supported chains, outputs, interface type, species coverage, and intended applications. This explorer operationalizes the UnivAIRRse framework by enabling rapid comparison and targeted identification of AIRR simulation tools.

##### 4.4.1 Adoption Trends and Research Networks

In conjunction with the interactive map, we analyzed publication trends and research networks in AIRR simulation studies from 2015 to 2025. Using VOSviewer, we constructed a keyword co-occurrence network, where nodes represent author keywords and links denote their concurrent presence in publications. Distinct clusters are identified by colors and qualitatively labeled based on predominant thematic terms (detailed methodology and clustering logic are provided in the Supplementary Section S5).

The bibliometric survey identified three principal thematic clusters, each distinctly color-coded in the co-occurrence network (Figure 4). The first cluster (green) represents foundational research focused on repertoire diversity and probabilistic generation models, including theoretical studies of generation probabilities, selection pressures, and sequence-level analyses of immunoglobulin repertoires. The second and largest cluster (red) corresponds to the clinical and translational domain, encompassing studies of immune responses, disease-associated repertoires (e.g., cancer), and immunotherapy, including antigen-driven investigations linking *in vivo* experimental systems with therapeutic design. The third cluster (blue) forms a molecular bridge, highlighting studies that characterize TCR-mediated recognition mechanisms and their structural interaction landscapes.

**Figure 4.**
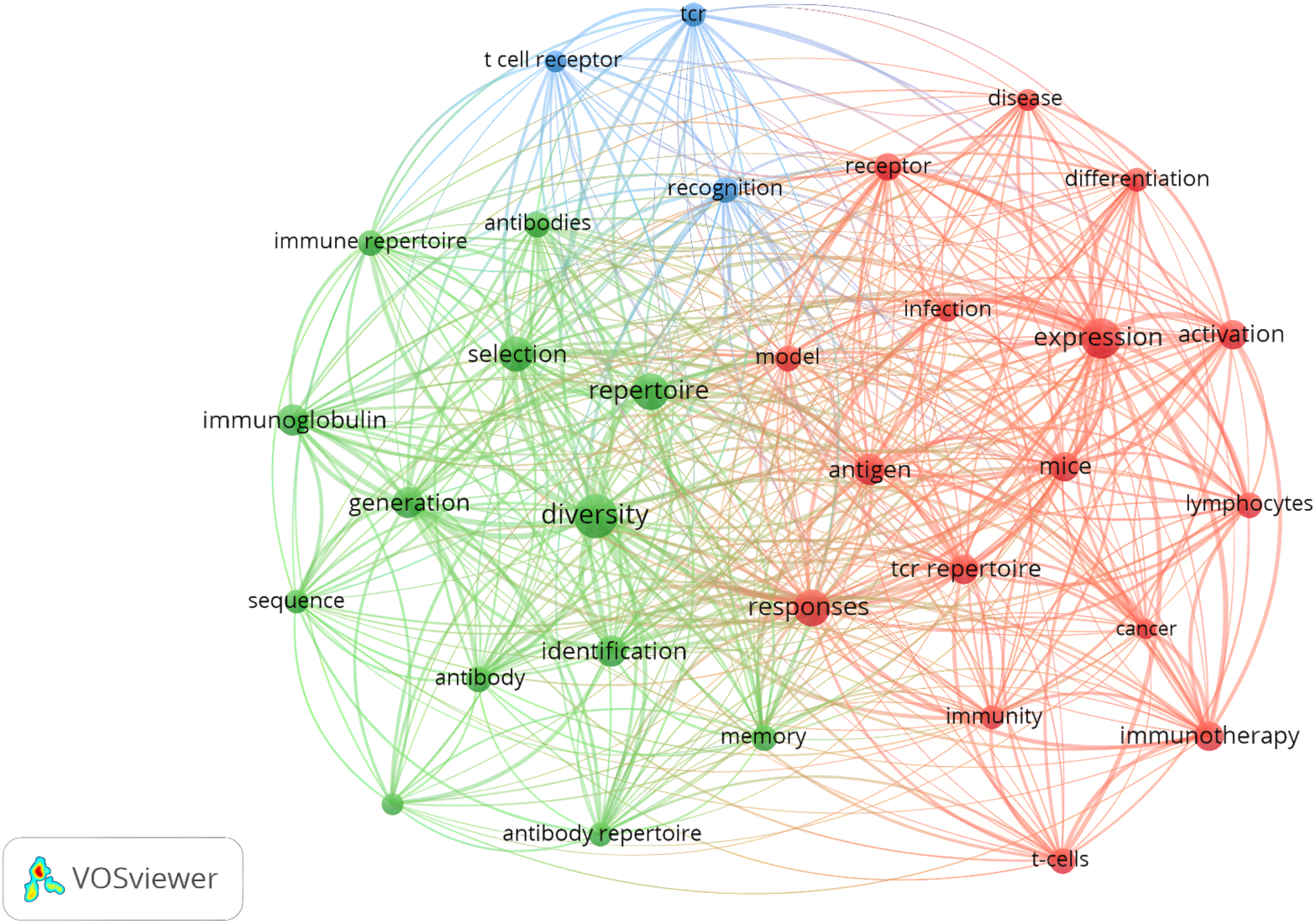
Keyword co-occurrence network of AIRR simulation research (2015–2025). In this network, nodes represent author keywords, and edges indicate their co-occurrence within publications. Colors denote clusters identified using the VOSviewer community-detection algorithm, revealing three primary thematic areas : (i) repertoire diversity and generative probabilities (green), (ii) TCR recognition and structural specificity (blue), and (iii) clinical responses and immunotherapy (red). Highly connected keywords such as model, response, and antigen occupy central network positions, acting as conceptual bridges between clusters and highlighting the role of predictive modeling as a unifying element linking theoretical, computational, and translational research domains within AIRR simulation.

Within this network, central bridging concepts, namely *model*, *response*, *antigen*, and *recognition*, occupy highly connected positions at the intersection of the three clusters. Their prominent connectivity indicates that theoretical generation models (green) and molecular recognition principles (blue) are increasingly integrated into translational research workflows (red). Collectively, this network topology suggests that AIRR simulation research has evolved into a cohesive and interconnected ecosystem in which foundational modeling, molecular recognition studies, and clinical applications increasingly inform one another.

The keyword co-occurrence network illustrates a progressive transition from theoretical and HMM-based frameworks in the initial phase to probabilistic generative models, such as IGoR and OLGA. More recently, this shift has extended to deep-learning and multimodal prediction tools, including TAPIR and TULIP, which integrate receptor sequences with transcriptomic and clinical metadata. The analysis of citation trends (Supplementary 5.8) corroborates this transition : established simulators exhibit stable and sustained usage, whereas newer machine-learning frameworks demonstrate rapid and ongoing growth. This evolution towards increasingly predictive and integrative tools aligns with the iterative, problem-driven logic of the UnivAIRRse framework, which is designed to connect foundational modeling approaches with advanced applications and to guide future innovations in computational immunology.

### 5. Use Cases and Adoption in Literature

Throughout this section, it is important to note that several categorizations and labels are derived from frequency of occurrence in the literature, reflecting prevailing usage rather than normative judgments. In various disciplines including immunology, computational biology, and applied medical studies, tools for simulating adaptive immune receptor repertoires have emerged as fundamental elements of investigative approaches. These systems offer more than just the creation of artificial data : they establish stable, repeatable settings that allow researchers to test data-processing workflows [26, 27]. In addition, they support the training and optimization of machine-learning and deep-learning models [28, 29]. Moreover, they facilitate the examination of natural processes that are difficult to extract from empirical collections [30, 31]. Such applications highlight the essential nature of these methods in validating pipelines, testing hypotheses, deriving insights into underlying biological principles, and advancing therapeutic applications [32–34].

Furthermore, these approaches are essential rather than optional. Key aspects of immune functions often elude detection within laboratory-derived sequencing information on adaptive immune receptors. For instance, authentic lineage tracing of cell clones, patterns in selection within antibody maturation sites, mechanisms for forming binding structures, and comprehensive profiles of antigen targets remain inaccessible [31, 35, 36]. Even extensive sequencing fails to uncover such hidden elements. Therefore, simulated frameworks provide the most reliable source of ground truth. As a result, these models address the divide between direct observations and required deductions, supporting thorough assessments of inference methods, interpretive instruments, and hypotheses about the immune system operational dynamics [37, 38].

#### 5.1 Benchmarking Analytical Tools

Simulators such as immuneSIM [10], AIRRSHIP [20], and Echidna [23]provide ground-truth labels for V(D)J assignment, clonotyping, lineage reconstruction, and single-cell integration [10, 12, 23, 39]. Across these studies, the authors consistently report high concordance for V and J gene calls but persistent ambiguity for D-gene assignment, as well as substantial variability in CDR3 annotations across pipelines [10, 40]. Single-cell simulators including Echidna also reveal limitations in multi-omic integration and transcriptome pairing [23]. In this study, we synthesize findings from previously conducted benchmarking studies without reanalyzing the original datasets. Table 2 presents a structured overview of the relevant benchmarking papers, including their tasks, truth labels, metrics, and key qualitative findings. The complete quantitative results and raw benchmarking data are accessible in the original publications.

**Table 2.**
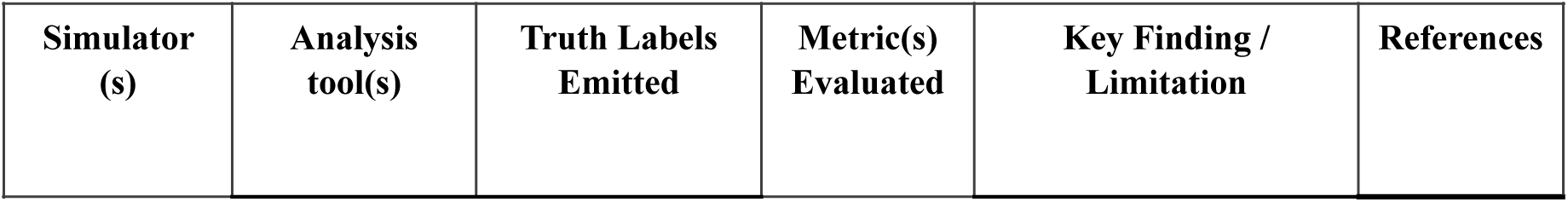
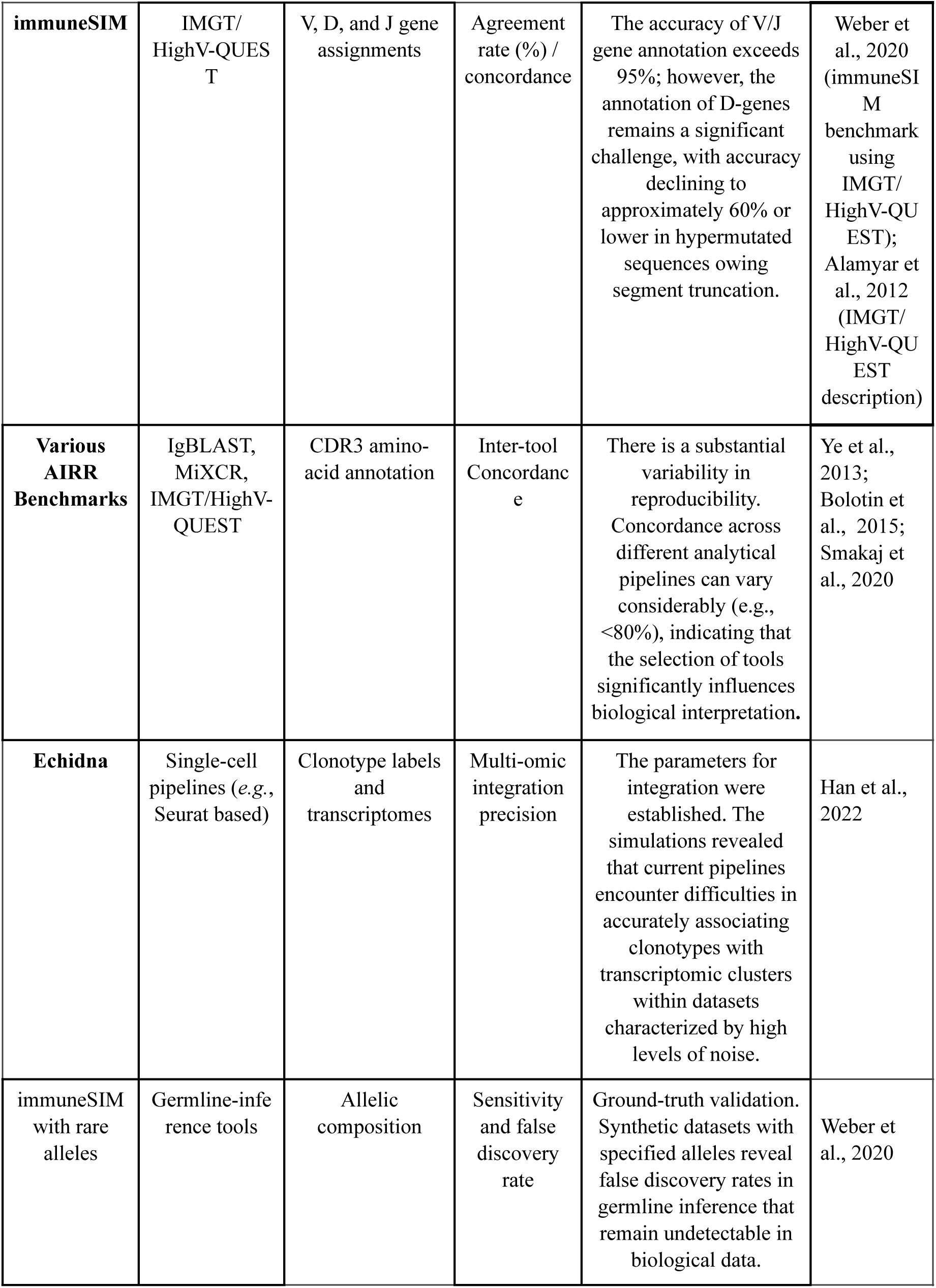
Summary of benchmarking studies in which AIRR simulators provide ground-truth labels for evaluating repertoire-analysis tools. It details the simulators and analysis pipelines employed, the characteristics of the truth labels produced, the metrics reported, and the key findings or limitations identified by each study. Detailed quantitative results and raw benchmarking data can be found in the original articles referenced.

| Simulator (s) | Analysis tool(s) | Truth Labels Emitted | Metric(s) Evaluated | Key Finding / Limitation | References |
| --- | --- | --- | --- | --- | --- |
| **immuneSIM** | IMGT/<br>HighV-QUEST | V, D, and J gene<br>assignments | Agreement<br>rate (%) /<br>concordance | The accuracy of V/J<br>gene annotation exceeds<br>95%; however, the<br>annotation of D-genes<br>remains a significant<br>challenge, with accuracy<br>declining to<br>approximately 60% or<br>lower in hypermutated<br>sequences owing<br>segment truncation. | Weber et<br>al., 2020<br>(immuneSIM<br>benchmark<br>using<br>IMGT/<br>HighV-QUEST);<br>Alamyar et<br>al., 2012<br>(IMGT/<br>HighV-QUEST<br>description) |
| **Various**<br>**AIRR**<br>**Benchmarks** | IgBLAST,<br>MiXCR,<br>IMGT/HighV-QUEST | CDR3 amino-<br>acid annotation | Inter-tool<br>Concordance | There is a substantial<br>variability in<br>reproducibility.<br>Concordance across<br>different analytical<br>pipelines can vary<br>considerably (e.g.,<br><80%), indicating that<br>the selection of tools<br>significantly influences<br>biological interpretation. | Ye et al.,<br>2013;<br>Bolotin et<br>al., 2015;<br>Smakaj et<br>al., 2020 |
| **Echidna** | Single-cell<br>pipelines (e.g.,<br>Seurat based) | Clonotype labels<br>and<br>transcriptomes | Multi-omic<br>integration<br>precision | The parameters for<br>integration were<br>established. The<br>simulations revealed<br>that current pipelines<br>encounter difficulties in<br>accurately associating<br>clonotypes with<br>transcriptomic clusters<br>within datasets<br>characterized by high<br>levels of noise. | Han et al.,<br>2022 |
| immuneSIM<br>with rare<br>alleles | Germline-inference<br>tools | Allelic<br>composition | Sensitivity<br>and false<br>discovery<br>rate | Ground-truth validation.<br>Synthetic datasets with<br>specified alleles reveal<br>false discovery rates in<br>germline inference that<br>remain undetectable in<br>biological data. | Weber et<br>al., 2020 |

Representative examples showing how simulators support evaluation of repertoire-analysis pipelines by providing defined ground-truth labels and quantitative metrics. These cases illustrate differences in accuracy, reproducibility, and sensitivity across gene assignment, clonotyping, and single-cell integration workflows.

#### 5.2 Validating Machine Learning Models and Clinical Pipelines

Large labeled AIRR-seq cohorts remain scarce, making simulation essential for training and validating machine-learning models. Simulators provide synthetic cohorts with controlled variation in sequencing depth, sampling bias, and noise, enabling robust calibration of ROC and precision–recall curves, rare-clone detection, trajectory inference for clinical monitoring, and evaluation of batch effects. These controlled settings prevent shortcut learning and help ensure that model performance reflects biological signals rather than dataset artifacts [30, 33, 41].

#### 5.3 Validating Biological Mechanisms

Mechanistic simulators replicate fundamental immunological processes, including recombination, somatic hypermutation, clonal expansion, and post-recombination selection. Tools such as IGoR [7], repgenHMM [8] and OLGA [16] allow controlled perturbation of recombination probabilities, mutation rates, and selection strengths. This enables systematic testing of how generative diversity, mutation clustering and selection pressures influence repertoire architecture [7, 8, 42].

To represent the immune system’s emerging complexity, advanced agent-based platforms recreate the shifting conditions within germinal centers and other lymphoid structures. Recent germinal center simulators, including C-ImmSim [43] and newer computational systems, merge processes across molecular, cellular, and tissue scales, allowing them to model how antigens, cytokine networks, and expanding clones influence one another. By using this integrated perspective, these tools reproduce core immune phenomena such as affinity maturation, memory formation, and the competitive dynamics that shape clonal selection and dominance [44, 45]. Ultimately, this modeling strategy provides a powerful means to explore how spatial organization and localized interactions collectively determine the trajectory of the broader immune response [43, 45].

##### Box 3

Hallmarks of Successful AIRR Simulators

Successful simulators share four defining traits:

1. **AIRR-format compatibility**, ensuring interoperability with community data standards.
2. **Modular design with transparent parameters**, allowing re-estimation and component replacement.
3. **Realism beyond marginal statistics**, capturing higher-order repertoire structures and cross-cohort generalization.
4. **Adherence to community standards**, facilitating validation and reuse across studies.

These qualities, not novelty alone, predict sustained adoption and long-term impact within the AIRR simulation ecosystem.

## 6. Limitations in Current AIRR Simulators

Despite major progress in modeling adaptive immune receptor repertoires, current AIRR simulators remain constrained by conceptual, biological, and computational limitations. These limitations arise from incomplete representation of key immunological processes, loss of contextual information during sequencing, insufficient modularity, and restricted interoperability with analytical pipelines. As a result, existing frameworks deviate from biological realism and do not fully capture the multi-scale and context-dependent nature of repertoire evolution across cellular, tissue, and individual levels.

A central limitation is that AIRR-seq data provide only a narrow molecular projection of a far more complex immunological state. Although sequencing preserves low-level properties such as V/J usage, CDR3 structure, somatic-hypermutation footprints, and UMI-normalized abundances [4, 15], it omits spatial organization, tissue residency, cytokine-signaling history, migration dynamics, and antigen-exposure chronology [14]. In practice, sequencing acts as a measurement operator that compresses a dynamic, multi-scale biological system into a static molecular snapshot. This projection is inherently lossy, shaped by sampling biases, tissue-specific recovery, PCR noise, and experimental dropout, all of which distort the relationship between underlying biology and observed repertoires [46].

Because of this lossy nature, simulators validated solely against marginal sequence statistics may generate repertoires that look realistic but fail to capture causal and mechanistic structure at the level of clonal dynamics, lineage topology, or repertoire overlap [2, 10]. Many frameworks also embed implicit and often undocumented assumptions when linking molecular sequences to cellular or tissue processes, which complicates cross-tool comparison and reduces reproducibility. Monolithic design remains another obstacle: several widely used simulators bundle recombination, mutation, selection, and sampling into a single rigid block, making it difficult to update components, incorporate new biology, or integrate with external pipelines.

A further challenge is the trade-off between biological realism and computational scalability. Mechanistic models that incorporate germinal-center dynamics or antigen-driven selection are computationally expensive, whereas scalable models tend to oversimplify affinity maturation, spatial heterogeneity, or tissue-specific activation. Together, these limitations underscore the need for next-generation simulation frameworks that incorporate explicit biological context, adopt modular architectures, and provide multi-scale ground truth to support rigorous benchmarking and digital-twin-ready immune modeling.

Table S12 summarizes the dominant gaps in current simulators, links them to representative tools, and outlines potential strategies for advancing biological fidelity and interoperability. Figure 5 complements the Table by mapping these gaps onto the B-cell developmental trajectory, highlighting which stages are well represented, partially modeled, or largely absent in current simulation frameworks.

**Figure 5.**
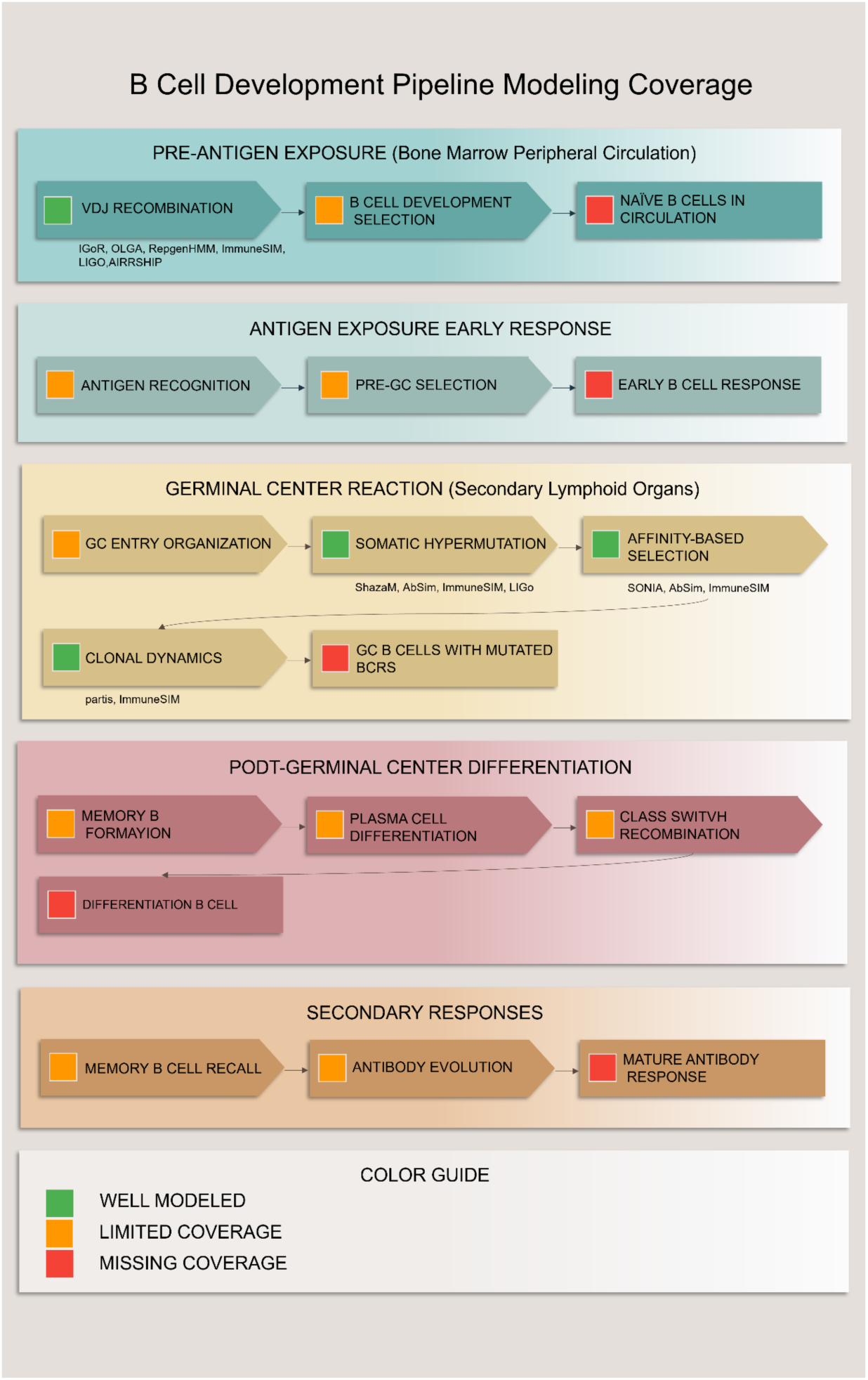
Biological coverage across the B-cell developmental trajectory. The figure illustrates how current AIRR simulators represent major stages of B-cell development, from V(D)J recombination and naïve repertoire formation to germinal-center evolution and post-GC differentiation. Green segments indicate processes that are well captured by existing tools (e.g., recombination, SHM patterns). Orange denotes partially modeled stages such as early clonal expansion, selection dynamics, and basic affinity maturation. Red highlights processes that remain largely absent, including cytokine-mediated feedback, spatial zonation within germinal centers, tissue trafficking, and secondary immune responses. The visualization underscores a biological gradient: phenomena closely tied to molecular readouts are reproduced with higher fidelity, whereas context-dependent, spatial, and temporal processes remain underrepresented.

### 6.1 Sequencing as a Molecular Proxy: What Is Observed and What Is Missing

AIRR sequencing offers a high-resolution molecular view of immune receptors but captures only a fraction of the full immunological context. While V/J usage, CDR3 composition, SHM patterns, and clonal abundance can be reliably inferred [4, 15], higher-order biological layers remain inaccessible. Spatial tissue architecture, migration paths, cytokine signaling history, antigen-exposure timing, and stochastic activation events are not encoded in the sequence record [14]. Consequently, AIRR-seq compresses a dynamic, multi-layered biological ecosystem into a reduced molecular representation, affected by sampling stochasticity, recovery bias, PCR noise, and dropout effects [46].

This lossy projection undermines validation strategies that rely only on sequence-level comparisons. Simulators tuned to match marginal sequence statistics may fail to reproduce higher-order structures such as lineage branching, clonal competition, or repertoire-overlap patterns [2, 10]. Temporal information is similarly absent from isolated sequences; it emerges only when sequences are placed within explicitly modeled clonal frameworks [19]. Despite this, some simulators assign immunological states (naïve, memory, activated) directly to synthetic sequences, even though such labels cannot be inferred from the variable region of sequences produced by AIRR-seq alone.

These constraints highlight that sequencing offers only a partial view of immune dynamics, underscoring the need for simulators that incorporate biological context beyond what AIRR-seq alone can reveal.

### 6.2 Formal Requirements for Context-Aware Simulation

Developing next-generation AIRR simulators that support rigorous biological inference requires adherence to principles of modularity, transparency, interoperability, and reproducibility.

A modular and hierarchical architecture is essential. Each biological process such as recombination, SHM, selection dynamics, and sampling mechanisms should be implemented as an independent but interoperable unit. Such modularity enables updates, empirical calibration, and controlled replacement of components without re-implementing the entire simulator.

Transparency and provenance are critical for reproducibility. Parameters, priors, default settings, model assumptions, and random seeds must be explicitly declared. Outputs should comply with AIRR-C standards and include metadata detailing parameterization, versioning, and computational provenance. Deterministic execution modes, containerized environments, and continuous-integration testing further ensure reproducible behavior across computational platforms.

Robust benchmarking requires multi-scale ground truth. Simulators must provide clonal labels, lineage trees, mutation histories, and repertoire level summaries, not just sequences, to support evaluation at sequence, clonal, repertoire, and population scales.

Integration and usability are equally important. Standardized APIs, extensive documentation, and command-line tools facilitate incorporation into machine-learning pipelines and multi-omics frameworks. Community registries, shared parameter sets, and coordinated benchmarking efforts are necessary to harmonize validation and situate simulators within emerging digital-twin ecosystems.

### 6.3 Summary of Gaps and Context Dependence

Current AIRR simulators successfully reproduce several molecular properties, including SHM patterns and diversity metrics, and have become indispensable tools in computational immunology. However, their realism remains strongly context-dependent. Four major gaps dominate:

1. Sequencing is frequently treated as a comprehensive representation of immune receptor biology. However, from a systems-level perspective, it represents only a partial and lossy projection that omits spatial and temporal context, activation history, and antigen exposure.
2. Implicit biological assumptions often remain undocumented. Links between cellular, tissue, and molecular layers are embedded within code rather than explicitly declared, reducing interpretability and cross-tool compatibility.
3. Monolithic architectures hinder extensibility. Many simulators combine multiple biological processes into rigid pipelines that cannot easily incorporate new biological knowledge or integrate with external systems.
4. Realism and scalability remain in tension. Highly detailed models are computationally expensive, whereas scalable models often oversimplify essential immunological dynamics.

Addressing these gaps requires explicit encoding of biological context, modular architectures, transparent assumptions, and unified validation frameworks that evaluate fidelity across sequence, clonal, repertoire, and population scales. Table ***S12*** summarizes these structural gaps, and Figure 5 illustrates how they map onto the B-cell developmental trajectory. To operationalize these conceptual limitations, Table ***S12*** distills them into modular categories that directly map onto the biological, computational and representational layers of the UnivAIRRse framework.

Together, Table S12 and Figure 5 demonstrate a clear gradient : processes directly tied to sequence-level readouts are modeled with higher fidelity, whereas context-dependent, spatial, and temporal phenomena remain underrepresented. Closing this gap will require modular, multi-scale simulators that encode biological context explicitly and can be validated across multiple data modalities. The next section outlines strategies for developing such biologically faithful and interoperable frameworks.

## 7. Future Directions : Toward Digital Twin–Ready Immune Simulation

The trajectory of AIRR simulation is increasingly converging with the broader vision of biomedical digital twins. In contemporary computational medicine, a digital twin is defined as a continuously updated, data-driven computational representation of an individual’s biological system, tightly coupled to multimodal and longitudinal observations [47, 48]. Unlike conventional simulators, which operate as static and unidirectional models, digital twins maintain a bidirectional, iteratively updated link with the patient. Model predictions guide sampling and intervention strategies, while new clinical data iteratively refine the internal model state.

Developing an immune digital twin requires integration across levels of biological organization. For adaptive immunity, this entails linking AIRR-seq data with cytokine landscapes, single-cell transcriptomes, antigen-specificity assays, clinical phenotypes, and potentially spatial tissue measurements [29, 49]. Such integration would enable continuous tracking of clonal trajectories during infection or vaccination, monitoring the emergence of protective or pathogenic clones, forecasting transitions between naïve, effector, and memory states, and supporting individualized therapeutic strategies.

Current AIRR simulators already provide foundational components for this vision. Tools such as IGoR, repgenHMM, immuneSIM, Echidna, and AIRRSHIP offer mechanistic models of recombination, somatic hypermutation, clonal expansion, and sequencing assays [2, 7, 10]. However, they are not yet digital twins. They lack personalized calibration based on patient-specific genomic and immunological characteristics, real-time assimilation of longitudinal data streams, multi-modal inference engines capable of integrating scRNA-seq, antigen mapping, cellular state transitions, and clinical metadata, as well as prospective clinical validation—each of which is essential for digital twin systems [49, 50].

Accordingly, current AIRR simulators should be conceptualized as *digital-twin–enabling modules* rather than complete digital twins. The path forward requires modular simulator architectures, flexible enough to incorporate biological processes as independent components; real-time updating frameworks that combine filtering algorithms, Bayesian inference, or neural differential equations; multi-omics integration pipelines capable of linking receptor sequences to cell states and clinical phenotypes; and uncertainty quantification methods that support robust prediction under heterogeneous data sources [51, 52].

When these components are unified within adaptive and dynamically calibrated frameworks, AIRR-based immune models can evolve into actionable clinical systems. Such digital twin–ready architectures could anticipate vaccine response trajectories, detect early signatures of immune dysregulation, optimize individualized immunotherapies, or predict relapse in autoimmune conditions. Ultimately, the transition from static simulation to continuous, personalized, and clinically integrated digital twin platforms represents the next major conceptual and technological frontier for computational immunology.

## 8. Conclusion

By positioning existing AIRR simulators within the UnivAIRRse framework, this review provides a unified structure for comparing their biological scope, abstraction level, and underlying assumptions. Sequence-level engines such as IGoR, OLGA, and repgenHMM operate closest to the Sequesphere. Repertoire-level frameworks including immuneSIM, AIRRSHIP, and partis extend into the Clonosphere and Repertoire domains, while specificity-oriented predictors such as TAPIR and TULIP explore the Specifisphere. Population-level models introduced by Ruiz Ortega and Böttcher [21] characterize public and private clone distributions across individuals, and analytical tools such as sumrep support standardized benchmarking and reproducibility. Collectively, these tools have established simulation as a central methodological foundation of modern immunoinformatics.

As the first unified coordinate system for organizing AIRR simulators, the UnivAIRRse framework provides a coherent lens for interpreting heterogeneous modeling assumptions across biological and computational scales. By formalizing this structure, this review establishes a baseline for future benchmarking, methodological comparison, and the systematic design of next-generation simulators.

The bibliometric analysis conducted in this study reveals a mature and rapidly evolving intellectual landscape within the field of AIRR simulation. The identification of three core, interconnected clusters such as repertoire generation, antigen-specificity prediction, and clinical translation highlights a well-defined research pipeline that extends from fundamental biology to translational applications. This structured ecosystem underscores the field’s readiness for a unifying framework capable of integrating these disparate domains.

Our analysis also reveals a clear methodological transition : from earlier theoretical models to advanced machine-learning frameworks for predictive tasks. The UnivAIRRse framework is designed to support this shift by providing a structured methodology that links foundational discovery to advanced application. In doing so, it offers a consistent reference point for future development and a practical tool for the diverse communities driving progress in computational immunology. Despite this progress, important limitations remain. Most simulators rely on AIRR sequencing as a partial and lossy molecular proxy, with limited incorporation of spatial, temporal, or tissue-level context. The mapping between cellular dynamics and molecular readouts often remains underconstrained, while transparency, tunability, and interoperability vary widely across platforms. The lack of unified provenance tracking, comprehensive ground-truth datasets, and scalable architectures further restricts the development of robust and generalizable simulation frameworks.

Looking forward, the concept of digital-twin–ready immune simulation provides a coherent roadmap for the next decade. Realizing this vision will require integrating modular simulators with longitudinal clinical and multi-omic datasets, enabling real-time synchronization and parameter updating, incorporating explicit uncertainty quantification, and achieving prospective clinical validation. A digital twin represents a continuously updated virtual model that reflects real biological behavior and adapts as new data arrive. When aligned with the broader digital twin paradigm [51, 53, 54], such frameworks can transform simulation from a static modeling exercise into a dynamic, data-driven, individualized predictive system capable of forecasting immune trajectories, evaluating therapeutic interventions, and informing clinical decisions.

In conclusion, AIRR simulation has progressed from a conceptual niche to an essential foundation of computational immunology. The next generation of simulators, defined by transparency, interoperability, contextual realism, and clinical validation, will move the field from descriptive modeling toward mechanistic understanding and from prediction toward actionable decision support. Ultimately, these advances will enable personalized vaccination strategies, optimized immunotherapies, and precise immune monitoring at both individual and population scales.

## Acknowledgments

The authors gratefully acknowledge the valuable comments and insightful suggestions provided by the IMGT® team during the preparation and revision of this manuscript. We especially thank Véronique Giudicelli, Gaoussou Sanou, and Patrice Duroux for their thoughtful feedback and support, which significantly improved the clarity, conceptual framing, and overall quality of this work.

## Notes

https://www.imgt.org/AIRR-Simulator/

### Competing Interest Statement

The authors have declared no competing interest.

