## Supplementary Materials for "UnivAIRRse: A Unified Framework for Organizing and Comparing Adaptive Immune Receptor Repertoire Simulators"

**Affiliations**

#### Supplementary Appendix I — Historical Evolution of AIRR Simulators

This appendix outlines the conceptual and technological progression of adaptive immune receptor repertoire (AIRR) simulators, spanning early theoretical models to modern AI-enabled, multimodal frameworks. Table S1 provides a structured timeline of major milestones, illustrating how increasing biological realism and computational sophistication have shaped the field.

***Table S1.*** *Historical evolution of AIRR simulators from theoretical prototypes to AI-integrated frameworks. This Table summarizes the conceptual and technological progression of AIRR simulators from early theoretical and shape-space models (1970s–1980s) to modern AI-integrated and multimodal frameworks (2020s–present). Each generation is characterized by its representative models, hallmark features, key innovations, scientific impact, current status, and known limitations, illustrating how the field has advanced toward increasingly realistic, data-driven, and context-aware immune-repertoire simulation.*

| Generation | Years | Representative Models | Hallmark Features | Key Innovations | Impact on Field | Current Status | Limitations |
|---|---|---|---|---|---|---|---|
| **Proto-simulators (Conceptual)** | 1970s – 1980s | Perelson & Oster (1979); Jerne (1974); Stewart & Varela (1980); Farmer et al. (1986) | Theoretical network/ shape-space models; abstract receptor representations | Shape-space formalism; network dynamics; analytical immune models | Established foundations for computational immunology | Historical foundation | No sequence-level realism; no empirical validation |

| | | | | | | | |
|---|---|---|---|---|---|---|---|
| **First Generation (Abstract Repertoire)** | 1990s – 2000s | Celada & Seiden (1992); Perelson (1997); Seiden & Celada (1998) | Bit-string repertoires; agent-based and cellular-automata simulations | Stochastic cell interactions; multi-scale abstraction | Demonstrated simulation as a method | Framework influence persists | Limited realism; no empirical grounding |
| **Transitional Generation (Toward Data-Driven)** | ~2005–2015 | Early partis; custom VDJ scripts | Probabilistic repertoires; empirical VDJ usage | Linking theory to sequences; basic clone-size modeling | Enabled emergence of modern simulators | Legacy generation | Non-standardized; limited scalability |
| **Second Generation (Data-Driven)** | 2016 – 2020 | IGoR; OLGA; SONIA; immuneSIM; AbSim; partis | Large AIRR training; explicit recombination; clonal expansion; SHM modeling | Standardized benchmarking; open-source frameworks | Defined modern standards | Widely used | Limited multi-omics; simplified tissue context |
| **Third Generation (AI-Integrated, Multimodal)** | 2020s – present | immuneML; Echidna; TAPIR; TULIP; deep generative models | Multimodal simulation (AIRR + scRNA + antigen); deep learning | Personalized repertoires; context-aware simulation | Enabling digital-twin development | Rapid growth | Data-hungry; interpretability limits; no clinical validation |

### Supplementary Appendix II – The UnivAIRRse Framework: Formal Definitions and Projection Operators

This appendix provides the formal details underlying the UnivAIRRse framework introduced in the main text (§2.3). UnivAIRRse is defined as a hierarchical coordinate system that organizes immune receptor representations into five operational levels. These levels span a continuum from experimentally observed molecular records to theoretical generative potential, providing a principled structure for simulation design, benchmarking, and interpretation.

#### 1. Formal Definitions of the Five UnivAIRRse Levels

Level 1:The Sequesphere (S1)

Definition:
The Sequesphere contains atomic, per-sequence molecular records obtained from AIRR-seq or simulators that emulate experimental output.

Objects in S1 include:

- Nucleotide and amino-acid sequences
- V/J (and D, if applicable) gene assignments
- UMI counts / molecule counts
- Somatic hypermutation calls and mutation profiles
- Paired chain information (for scAIRR-seq or simulated pairing)
- Provenance metadata: sample ID, timepoint, read quality, cell barcode, assay information

Interpretation:
S1 is the closest representation to experimental truth; it carries concrete, observable information with well-characterized technical error models (PCR bias, sequencing errors, UMI noise).

Level 2:The Clonosphere (S2)

Definition:
The Clonosphere groups S1 sequences into clonal families derived from a single V(D)J rearrangement and its mutational descendants.

Each clonal object includes:

- Clonal identifiers
- Reconstructed lineage trees (topology + branch lengths)
- Mutation trajectories and SHM event logs
- Optional affinity/fitness annotations

Interpretation:
S2 introduces inference: lineage reconstruction, SHM models, and clonal assignment thresholds. Outputs are model-dependent and not directly observable.

Level 3:The Specifisphere ($S_3$)

Definition:
The Specifisphere groups receptors by functional antigen recognition, forming specificity neighborhoods.

Each object represents:

- A set of TCRs/BCRs recognizing the same antigen/epitope
- Functional similarity scores or embeddings
- Cross-reactivity structure
- Optional mechanistic/structural links (motifs, CDR3 features)

Interpretation:
$S_3$ is derived from functional assays, ML prediction, or structural modeling.
It is inherently inferential, as specificity cannot be identified from sequence alone.

Level 4:The Repertoire (S4)

Definition:
The Repertoire level is the set of all sequences biologically present in an individual (or sampled compartment) at a specific time t and sampling S.

Each repertoire includes:

- Global diversity indices (Shannon, Simpson)
- Clone-size distributions
- Isotype distributions (for BCRs)
- Population-level summaries
- Sampling design (sequencing depth, tissue, timepoint)

Interpretation:
S4 aggregates S1 sequences and S2 clonal families into system-level structures.

Level 5: The UnivAIRRse (S5)

Definition:
The UnivAIRRse is the space of all possible immune receptors generable by:

- Germline gene repertoires
- V(D)J recombination mechanisms
- Junctional diversity
- Somatic hypermutation pathways
- Evolutionary constraints

Interpretation:
S5 is a theoretical generative landscape. It defines what is possible, not what is observed.

#### 2. Projection Operators Between Levels

Each pair of adjacent levels is connected by a projection operator that transforms one representation into another. These operators structure how simulators should specify their assumptions.

##### P1→2 : Sequesphere → Clonosphere

Groups individual sequences into clones.
Depends on:

- Clonal assignment rule (e.g., CDR3 identity, V/J similarity, probabilistic ancestry)
- Phylogenetic inference model
- SHM model and thresholds

##### P2→3 : Clonosphere → Specifisphere

Maps clones to antigen-specific neighborhoods.
Depends on:

- Binding assays
- ML-based specificity predictions
- Epitope similarity models
- Structural docking / motif constraints

##### P3→4 : Specifisphere → Repertoire

Constructs system-level composition.
Depends on:

- Expansion/contraction dynamics
- Organ/tissue sampling
- Clonal competition
- Repertoire turnover

##### P4→5 : Repertoire → UnivAIRRse

Projects observed/system-level repertoires into the theoretical generative space.
Depends on:

- Germline library
- Insertion/deletion models
- Probabilistic generative models (Pgen)
- Mutational space reachable via SHM

These operators allow simulators to declare which biological processes they model and where simplifying assumptions enter.

#### 3. Temporal Semantics in UnivAIRRse

Time is treated differently at each level:

| Level | Temporal Meaning |
|---|---|
| S1 Sequesphere | Timeless* single-molecule observations at sample time t |
| S2 Clonosphere | Temporal dynamics encoded via lineage tree topology & branch lengths |
| S3 Specifisphere | Temporal changes reflect expansion/contraction of specificity neighborhoods |
| S4 Repertoire | Longitudinal dynamics via shifts in repertoire composition |
| S5 UnivAIRRse | Timeless combinatorial possibility space |

Key principle*:
Time is only biologically meaningful in the Clonosphere and above. Sequence-level data restricted to the variable region (V(D)J) cannot identify temporal labels such as naïve, primary, or memory states. Only when constant-region genes (C genes) or class-switch information are present can we infer partial temporal ordering, but this is still insufficient for full biological time reconstruction.

#### 4. Empirical–Theoretical Gradient

The UnivAIRRse hierarchy defines an epistemological gradient:

- Concrete → Abstract
- Observable → Inferred → Theoretical
- Experiment-proximal → Model-dependent → Hypothetical

| Level | Observability | Dependence on Assumptions |
|---|---|---|
| S1 | Directly observable | Low |
| S2 | Partially observable (requires inference) | Moderate |
| S3 | Indirect / functional inference | High |
| S4 | Statistical aggregation | Medium |
| S5 | Purely theoretical | Very high |

This clarifies what simulators can validate and what must remain assumption-driven.

**5. Purpose of the UnivAIRRse Framework in Simulation**

UnivAIRRse provides:

<u>5.1 A Coordinate System for Comparing Simulators</u>

Simulators can now be classified by:

- Which levels they produce
- Which projection operators they model
- Which assumptions they implement explicitly or implicitly

<u>5.2 A Transparency Tool</u>

We recommend that developers clearly indicate:

- Where inference vs. observation occurs
- Which biological processes are approximated
- Which outputs are ground truth

<u>5.3 A Benchmarking Standard</u>

Evaluations are most meaningful when aligned with the appropriate level:

- Sequence realism → S1
- Clonal structure accuracy → S2
- Specificity predictions → $S_3$
- Diversity and systemic properties → S4
- Generative validity → S5

#### Supplementary Appendix III — Benchmarking, Systemic Limitations, and Digital-Twin Requirements

This appendix summarizes (i) how AIRR simulators support benchmarking of repertoire-analysis pipelines, (ii) systemic limitations in existing tools, and (iii) requirements for next-generation digital-twin immune simulators.

***Table S2.*** *Categorization of AIRR simulators by abstraction level*

*This table classifies AIRR simulators according to the level of abstraction at which they model immune processes. The categories range from nucleotide-level recombination mechanisms to cohort-scale population statistics. Each abstraction layer reflects specific modeling priorities, typical ground-truth labels, and characteristic use cases.*

| **Abstraction Level** | **Representative Tools (Stable IDs)** | **Primary UnivAIRRse Domain(s)** | **Typical Outputs / Ground Truth** | **Example Use Cases** |
|---|---|---|---|---|
| **Molecular level** | repgenHMM (1), IGoR (2), OLGA (3) | Sequesphere | Rearranged sequences; generation probabilities (Pgen); recombination event logs | Modeling V(D)J recombination; generating naïve repertoires; priors for selection models |
| **Clonal level** | partis (11), AIRRSHIP (10), AbSim (8), Echidna (9), SHM models (10a) | Clonosphere | Clonal identifiers; lineage trees; SHM trajectories; per-cell clone identities (single-cell) | Benchmarking clonotyping and lineage inference; modeling SHM and affinity maturation |
| **Repertoire level** | immuneSIM (7), SONIA (4), soNNia (5), sumrep (12) | Repertoire | Clone-size distributions; diversity indices; post-selection probabilities (Ppost); summary statistics | Comparing simulated vs. real repertoires; evaluating selection pressures; QC and reproducibility |

| | | | | |
|---|---|---|---|---|
| **Functional level** | TAPIR (13), TULIP (14) | Specifisphere | Predicted receptor–antigen scores; binding probabilities; functional signatures | Epitope prediction; vaccine design; translational immunotherapy |
| **Population level** | Ruiz Ortega model (15), Böttcher model (16) | Population level | Publicness metrics; repertoire-sharing probabilities; cohort-level variation | Analyzing public vs. private clones; cross-individual comparisons |
| **Measurement layer** | InSilicoSeq 2.0 (17) | Measurement layer | Synthetic reads with known error positions; sequencing-platform bias profiles | Testing pipeline robustness; UMI/PCR-bias calibration; end-to-end sequencing evaluation |

***Table S3.*** *Categorization of AIRR simulators by application focus*

*This Table classifies representative AIRR simulators according to their primary application domain, including benchmarking, personalization, translational prediction, and educational or hypothesis-testing use. Although many tools can support multiple applications, one dominant purpose typically guides their design, implementation, and adoption.*

| Application Focus | Representative Tools / Models | Primary Use Cases |
|---|---|---|
| **Benchmarking-focused (Clonal, Repertoire, Measurement)** | immuneSIM (6); AIRRSHIP (8); sumrep (12); AbSim (7); Echidna (9); InSilicoSeq 2.0 (17) | Evaluation of clonotyping and phylogenetic methods; assessment of repertoire-diversity metrics; validation of single-cell integration workflows; testing of sequencing-error and assay models |
| **Personalization-focused (Sequesphere, Repertoire, Population)** | repgenHMM (1); IGoR (2); OLGA (3); SONIA (4); soNNia (5) | Modeling individualized recombination processes; generating subject-specific repertoires; reconstructing personalized selection landscapes |

| | | |
|---|---|---|
| **Prediction and translational modeling (Specifisphere, Population)** | TAPIR (13); TULIP (14); Ruiz Ortega model (15); Böttcher model (16) | Antigen/epitope prediction; modeling public/private clones; identifying disease-associated repertoire patterns; supporting immunotherapy and vaccine design |
| **Educational / hypothesis-testing frameworks (Clonal, Repertoire)** | immuneSIM (6); partis (paired) (11) | Teaching repertoire dynamics; illustrating lineage reconstruction; enabling exploratory simulations for conceptual or qualitative hypothesis generation |

***Table S4.*** *Benchmarking performance of representative AIRR-analysis tools using simulator-derived ground truth. This Table summarizes benchmarking studies in which AIRR simulators or synthetic repertoires were used to evaluate the accuracy, reproducibility and robustness of widely used repertoire-analysis pipelines. For each tool, the table lists the available truth labels, the metrics applied for evaluation and the principal findings. The results reveal major sources of variability including D-gene ambiguity, inconsistent CDR3 annotation and challenges in multi-omic integration, underscoring the need for standardized benchmarking frameworks for reliable AIRR-seq analysis.*

| Simulator | Truth Labels | Metric(s) Evaluated | Key Finding |
|---|---|---|---|
| **immuneSIM + IMGT/HighV-QUEST** | V/J assignments | Concordance | >97% accuracy for V/J; ~60% for D-gene → persistent ambiguity |
| **IgBLAST, MiXCR** | CDR3 amino-acid annotations | Reproducibility | 4.3–77.6% variability → need for standardization |
| **Echidna** | Clonotype + transcriptome pairs | Sensitivity; integration metrics | Reveals multi-omic integration challenges |
| **immuneSIM (rare alleles)** | Germline variants | Sensitivity/specificity | Enables robust evaluation of allele-discovery |

***Table S5.****Structural and methodological limitations of current AIRR simulators identified through gap analysis. This Table outlines major missing features and design constraints observed across widely used AIRR simulators. For each limitation, the representative tools and the resulting consequences for biological realism, inference accuracy and reproducibility are listed. The gaps include incomplete contextual modeling, hidden and non-tunable assumptions, insufficient provenance tracking, limited ground-truth availability, weak integration with multi-omics and AI workflows, scalability constraints and inadequate documentation. These structural issues highlight the need for modular, transparent and context-aware next-generation simulation frameworks.*

| Missing Feature | Representative Tools | Consequences |
|---|---|---|
| Sequencing treated as complete (no spatial/ temporal context) | partis, immuneSIM, AIRRSHIP | Cannot reproduce true lineage/tissue dynamics |
| Implicit, non-tunable biological assumptions | IGoR, OLGA, SONIA | Hidden biases; limited inference generality |
| Limited provenance tracking | Most tools | Reduced reproducibility |
| Partial or missing ground truth | immuneSIM (partial), others | Restricts lineage-level evaluation |
| Weak multi-omics/AI integration | SONIA, immuneSIM | Restricts translational modeling |
| Realism vs. scalability trade-offs | immuneSIM, lightweight simulators | Computational limits or oversimplification |
| Poor documentation/UX | Several tools | Low adoption |

***Table S6.*** *Core requirements for developing digital-twin–ready AIRR simulators. This Table summarizes the essential technological and biological capabilities needed for next-generation AIRR simulators that can support digital-twin applications. For each requirement, the corresponding functional features, existing gaps in current tools and the key opportunities for advancement are outlined. The requirements span personalization, continuous data integration, bidirectional modeling, calibrated validation, interoperability and scalable computation, highlighting the steps necessary for creating clinically actionable and dynamically updated immune digital twins.*

| Requirement | Features Needed | Current Gaps | Opportunity |
|---|---|---|---|
| Personalization | Per-patient calibration | Mostly population-level | Integrate clinical AIRR-seq and genotype |
| Continuous Data Integration | Time-series + spatial + scRNA input | No real-time updating | Streaming pipelines linked to EHR |
| Bidirectional Modeling | Intervention simulation | One-way generation | Predict therapy/ vaccine outcomes |
| Validation & Uncertainty | Calibrated predictions | Limited experimental validation | Prospective clinical benchmarking |
| Interoperability | Standardized APIs, FAIR | Fragmented formats | Full AIRR-C compliance |
| Scalability | Real-time simulation | HPC required | Cloud-native architectures |

#### Supplementary Appendix IV— Simulator Profiles, Feature Matrices, and UnivAIRRse Mapping

This appendix provides detailed descriptions of representative AIRR simulators, their functional capabilities, and their mapping within the UnivAIRRse hierarchy.

***Table S7.*** *Mini-profiles of representative AIRR simulators summarizing origin, core purpose, key functionalities and mapping within the UnivAIRRse hierarchy. This table provides a concise comparative overview of widely used AIRR simulators, including their year of introduction, development origin, principal modeling objective and defining features. Each entry specifies whether the tool supports BCR, TCR or both receptor classes and indicates the corresponding UnivAIRRse domain coverage. Together, these profiles highlight the diversity of modeling strategies across recombination inference, clonal evolution, repertoire-level simulation, specificity prediction, population-level modeling and sequencing-assay emulation.*

| # | Tool (Reference) | Year | Maintainer / Origin | Core Purpose | Key Features | BCR/ TCR | UnivAIRRse Levels |
|---|---|---|---|---|---|---|---|
| **1** | **repgenHMM** (Elhanati 2016) | 2016 | ENS Paris | Recombination inference | HMM-based VDJ model; generative sequences | Both | Sequesphere |
| **2** | **IGoR** (Marcou 2018) | 2018 | Yale | VDJ inference & generation | Probabilistic model; SHM for BCR | Both | Sequesphere, Repertoire |
| **3** | **OLGA** (Sethna 2019) | 2019 | Yale / ENS | Pgen computation | Fast motif sampling; publicness estimation | Both | Sequesphere |
| **4** | **SONIA** (Isacchini 2020) | 2020 | ENS Paris | Selection modeling | Post-generation filters; population variation | TCR | Repertoire, Population |
| **5** | **soNNia** (Isacchini 2021) | 2021 | ENS Paris | Deep-learning selection | Neural selection landscapes | Both | Repertoire, Population |
| **6** | **immuneSIM** (Weber 2020) | 2020 | ETH Zurich | Full repertoire simulation | SHM, clonality, motif implants, error model | Both | Clonosphere, Repertoire |
| **7** | **AbSim** (Yermanos 2017) | 2017 | ETH | Affinity maturation simulation | Lineage divergence; SHM cycles | BCR | Clonosphere, Repertoire |
| **8** | **AIRRSHIP** (Guest 2023) | 2023 | Oxford | BCR recombination simulator | Realistic junctions; recombination ground truth | BCR | Sequesphere, Clonosphere |
| **9** | **Echidna** (Yermanos 2023) | 2023 | ETH Zurich | Single-cell repertoire simulation | Paired AIRR + scRNA-seq | Both | Clonal, Repertoire |

| | | | | | | | |
|---|---|---|---|---|---|---|---|
| 10 | **SHM models** (Yaari 2013) | 2013 | Kleinstein Lab | SHM mutation modeling | Motif-based targeting & substitution | BCR | Clonosphere |
| 11 | **partis (paired)** (Ralph 2022) | 2022 | FHCRC | Paired-chain clonal inference | Heavy/light pairing + simulation | BCR | Clonosphere, Repertoire |
| 12 | **sumrep** (Olson 2019) | 2019 | Yale | Statistical evaluation | Summary metrics, repertoire divergence | Both | Repertoire |
| 13 | **TAPIR** (DeWitt 2023) | 2023 | Stanford | TCR–epitope DL prediction | CNN-based specificity modeling | TCR | Specifisphere |
| 14 | **TULIP** (Lu 2023) | 2023 | U. Washington | Unsupervised TCR–epitope mapping | Transformer architecture | TCR | Specifisphere |
| 15 | **Ruiz Ortega model** (2023) | 2023 | ENS | Public-clone statistics | Pgen×Ppost models | Both | Population |
| 16 | **Böttcher model** (2023) | 2023 | Germany | Analytical clone sharing | Public/private clone distributions | Both | Population |
| 17 | **InSilicoSeq 2.0** (Rösti 2023) | 2023 | Lausanne | Sequencing assay simulation | Read errors, platform-specific bias | DNA/ RNA | Measurement layer |

***Table S8.*** *Comprehensive feature and capability matrix for AIRR simulators across biological, computational and analytical dimensions.*

*This Table integrates the major functional attributes of representative AIRR simulators, covering biological realism, sequencing and error modeling, calibration flexibility, paired-chain support, ground-truth availability, personalization options, machine-learning and multi-omics integration, benchmarking utility and relevance to autoimmune studies. The matrix also specifies the UnivAIRRse domains represented by each tool. Collectively, this overview highlights key strengths, partial capabilities and missing functionalities across current AIRR-simulation frameworks.*

*Legend: ✓ = supported — = not supported • = partial*

| Tool | SHM | Error Model | Calibration | Paired Chains | Ground Truth | Personalization | ML/DL Integration | Multi-Omics | Benchmark Utility | Autoimmunity Relevance | UnivAIRRse Domains |
|---|---|---|---|---|---|---|---|---|---|---|---|
| **repgenHMM** | — | — | ✓ | — | — | ✓ | • | — | ✓ | ✓ | Sequesphere |
| **IGoR** | ✓(BCR) | — | ✓ | — | • | ✓ | • | — | ✓ | • | Sequesphere, Repertoire |
| **OLGA** | — | — | ✓ | — | — | ✓ (via IGoR) | • | — | ✓ | • | Sequesphere |
| **SONIA** | — | — | ✓ | — | — | ✓ | ✓ (basis of soNNia) | — | ✓ | ✓ | Repertoire, Population |
| **soNNia** | — | — | ✓ | — | — | ✓ | ✓✓ (deep learning) | — | ✓ | ✓ | Repertoire, Population |
| **immuneSIM** | ✓ | Optional | ✓ | — | ✓ | ✓ | ✓ (motif implants) | — | ✓✓ | ✓ | Clonosphere, Repertoire |
| **AbSim** | ✓ | — | ✓ | — | ✓ | • | — | — | ✓ | ✓ | Clonosphere, Repertoire |
| **AIRRSHIP** | — | — | ✓ | — | ✓ | • | — | — | ✓ | ✓ | Sequesphere, Clonosphere |
| **Echidna** | ✓ | — | ✓ | ✓ | ✓ | ✓ | ✓ | ✓✓ | ✓✓ | • | Clonal, Repertoire, Multi-omics |
| **SHM models** | ✓ | — | ✓ | — | — | • | • | — | ✓ | ✓ | Clonosphere |

| | | | | | | | | | | | |
|---|---|---|---|---|---|---|---|---|---|---|---|
| **partis (paired)** | ✓ | — | ✓ | ✓ | ✓ | ✓ | • | ✓ | ✓ | ✓ | Clonosphere, Repertoire |
| **sumrep** | N/A | N/A | ✓ | — | — | — | ✓ (stats for ML) | — | ✓✓ | • | Repertoire |
| **TAPIR** | — | — | ✓ | — | — | — | ✓✓ | — | ✓ | • | Specifisphere |
| **TULIP** | — | — | ✓ | — | — | — | ✓✓ (Transformer) | — | ✓ | • | Specifisphere |
| **Ruiz Ortega model** | — | — | ✓ | — | — | ✓ | ✓ (compatible) | — | ✓ | ✓ | Population |
| **Böttcher model** | — | — | ✓ | — | — | ✓ | • | — | ✓ | ✓ | Population |
| **InSilicoSeq 2.0** | N/A | ✓ | ✓ | — | ✓ (error truth) | — | • | — | ✓✓ | • | Measurement Layer |

***Table S9.*** *High-level categorization of AIRR simulation frameworks based on their primary modeling focus and defining features. This Table provides an overview of representative AIRR simulators grouped according to their dominant functional category, including sequence-level generators, selection models, repertoire-level frameworks, predictive specificity models, population-level statistical tools and sequencing-assay simulators. For each tool, the table lists the key modeling characteristics and the corresponding reference, offering a concise comparison of how different frameworks address distinct aspects of immune-repertoire simulation and analysis.*

| # | Tool | Category | Key Features | Reference |
|---|---|---|---|---|
| **1** | **repgenHMM** | Sequence-level | HMM inference of recombination; generative model | Elhanati 2016 |
| **2** | **IGoR** | Sequence-level | Probabilistic annotation; SHM (BCR) | Marcou 2018 |
| **3** | **OLGA** | Sequence-level | Fast Pgen; motif sampling | Sethna 2019 |

| | | | | |
|---|---|---|---|---|
| 4 | **SONIA** | Selection model | Post-generation selection; cohort variation | Isacchini 2020 |
| 5 | **soNNia** | Selection model (DL) | Neural selection landscapes | Isacchini 2021 |
| 6 | **immuneSIM** | Repertoire-level | SHM, clonality, motifs | Weber 2020 |
| 7 | **AbSim** | Repertoire-level | Affinity maturation cycles | Yermanos 2017 |
| 8 | **AIRRSHIP** | Repertoire-level | Realistic BCR junctions; ground truth | Guest 2023 |
| 9 | **Echidna** | Repertoire-level | Paired AIRR + scRNA | Yermanos 2023 |
| 10 | **SHM models** | Repertoire-level | Motif-based SHM | Yaari 2013 |
| 11 | **partis (paired)** | Repertoire-level | Paired-chain clonal inference | Ralph 2022 |
| 12 | **sumrep** | Evaluation utility | Summary metrics; divergence | Olson 2019 |
| 13 | **TAPIR** | Predictive (specificity) | DL receptor–epitope prediction | DeWitt 2023 |
| 14 | **TULIP** | Predictive (specificity) | Transformer epitope inference | Lu 2023 |
| 15 | **Ruiz Ortega model** | Population-level | Public/private clone statistics | Ruiz Ortega 2023 |
| 16 | **Böttcher model** | Population-level | Analytical clone-sharing | Böttcher 2023 |
| 17 | **InSilicoSeq 2.0** | Assay / Error simulation | Synthetic reads, error profiles | Rösti 2023 |

***Table S10.*** *Mapping of AIRR simulators to their corresponding UnivAIRRse domains based on supported representation levels. This Table shows how each AIRR simulator aligns with the five layers of the UnivAIRRse framework, indicating whether a tool supports sequence-level, clonal-level, repertoire-level, specificity-level or population-level representations. Tick marks identify full support, partial support or absence of functionality. This mapping highlights the conceptual scope and biological resolution of each simulator and clarifies how different tools occupy distinct regions of the immune-repertoire abstraction hierarchy.*

*Legend: ✓ = supported    — = not supported    • = partial*

| Tool | Seq | Clonal | Repertoire | Specificity | Population |
|---|---|---|---|---|---|
| **repgenHMM** | ✓ | — | — | — | — |
| **IGoR** | ✓ | — | • | — | — |
| **OLGA** | ✓ | — | — | — | — |
| **immuneSIM** | ✓ | ✓ | ✓ | • | — |
| **AIRRSHIP** | ✓ | ✓ | ✓ | — | — |
| **partis (paired)** | • | ✓ | ✓ | — | — |
| **SHM models** | ✓ | • | — | — | — |
| **SONIA** | — | — | ✓ | — | • |
| **soNNia** | — | — | ✓ | — | ✓ |
| **sumrep** | — | — | ✓ | — | • |
| **Ruiz Ortega model** | — | — | — | — | ✓ |
| **Böttcher model** | — | — | — | — | ✓ |
| **TAPIR** | — | — | — | ✓ | — |
| **TULIP** | — | — | — | ✓ | — |
| **Echidna** | ✓ | ✓ | ✓ | — | — |
| **InSilicoSeq 2.0** | — | — | — | — | — |

### Supplementary Appendix V— Bibliometric Analysis of AIRR Simulation Research

#### S5.1 Search Strategy and Data Extraction

A bibliometric analysis was conducted using the Web of Science database, concentrating on publications from 2015 to 2025. The search strategy was designed to target the topic field (TS; title, abstract, author keywords, Keywords Plus) with both proximity and phrase matching enabled. The query was constructed to identify literature at the intersection of immune repertoire terminology and computational modeling.

```
TS=(("immune repertoire" OR "adaptive immune repertoire" OR "lymphocyte repertoire" OR "antibody repertoire" OR "T-cell receptor repertoire" OR "TCR repertoire" OR "B-cell receptor repertoire" OR "BCR repertoire" OR "VDJ recombination" OR "immunoglobulin repertoire") AND (simulat* OR model* OR "in silico" OR comput* OR virtual OR "stochastic simulation" OR "agent-based model" OR "Monte Carlo simulation" OR "generative model"))
```

The analysis was restricted to English-language articles and reviews.

#### S5.2 Network Construction and Visualization Parameters

Bibliographic data were extracted and analyzed using VOSviewer (version 1.6.20) to construct keyword co-occurrence networks. This methodology elucidates the conceptual framework of a field by evaluating the frequency with which keywords co-occur within the same documents. To ensure clarity and reproducibility, the following configuration was used:

- **Keyword Source: Author keywords and Keywords Plus, as provided by the Web of Science.**
- **Thresholds:** To reduce extraneous noise and focus on well-established concepts, only keywords that appeared at least five times were included in the visualization.
- **Method of Analysis:** We employed the Full Counting technique to calculate link strength and utilized the Association Strength method for normalization.
- In visual representations, the size of a node corresponds to its frequency of occurrence, the thickness of an edge signifies the strength of co-occurrence, and the distance between nodes indicates their conceptual relationship.
- **Terminological Strategy:** We deliberately adopted a conservative approach with respect to the unification of synonyms. Specifically, we did not employ an automated thesaurus to consolidate biologically related terms (e.g., T-cell receptor vs. TCR; antibody vs. immunoglobulin). This decision enables the network to capture terminological heterogeneity, which is an inherent characteristic of this interdisciplinary field, where distinct sub-communities (e.g., computational vs. clinical) may prefer different vocabularies.

#### S5.3 Cluster Identification and Thematic Labeling

**Given that the clusters are generated through mathematical processes, specifically modularity maximization, and remain unlabeled by the software, the authors employed a qualitative approach to thematic naming. The assignment of cluster labels was executed through a structured interpretative process involving:**

- **Dominant Term Assessment:** Identification of keywords distinguished by high frequency and significant total link strength within each cluster (*e.g., immune repertoire, diversity, and generation*).
- **Conceptual Coherence:** An assessment of the extent to which groups of biologically related terms collectively represent shared research questions.
- **Addressing Terminological Overlap:** We explicitly acknowledge that several key terms (*e.g., receptor, model, response, and antigen*) encompass broad conceptual scopes. In the absence of enforced synonym integration, these terms function as bridging concepts that connect rather than delineate clusters.

Consequently, the cluster labels reflect the primary conceptual emphasis of each group rather than functioning as an exclusive classification. The redundancy observed among related terms is indicative of linguistic diversity within the field.

#### S5.4 Co-authorship Network Analysis

To offer a comprehensive perspective on the research landscape, we analyzed co-authorship networks. The co-authorship network (Figure S5) illustrates a well-established research community characterized by a hub-and-spoke configuration, which is indicative of effective knowledge dissemination and collaboration. The analysis identified distinct roles for prominent researchers (detailed metrics in Table S10): (i) Primary Network Hubs: Led by Victor Greiff and Sai T. Reddy, who emerged as central coordinators with the highest total link strength. (ii) Auxiliary Network Hubs: Aleksandra M. Walczak, Thierry Mora, and Alexander Yermanos form secondary hubs. The Walczak–Mora collaboration is particularly robust and anchors the statistical physics paradigm. (iii) Theoretical Leaders: Gur Yaari and Steven H. Kleinstein is highly cited, reflecting foundational contributions to the field's frameworks.

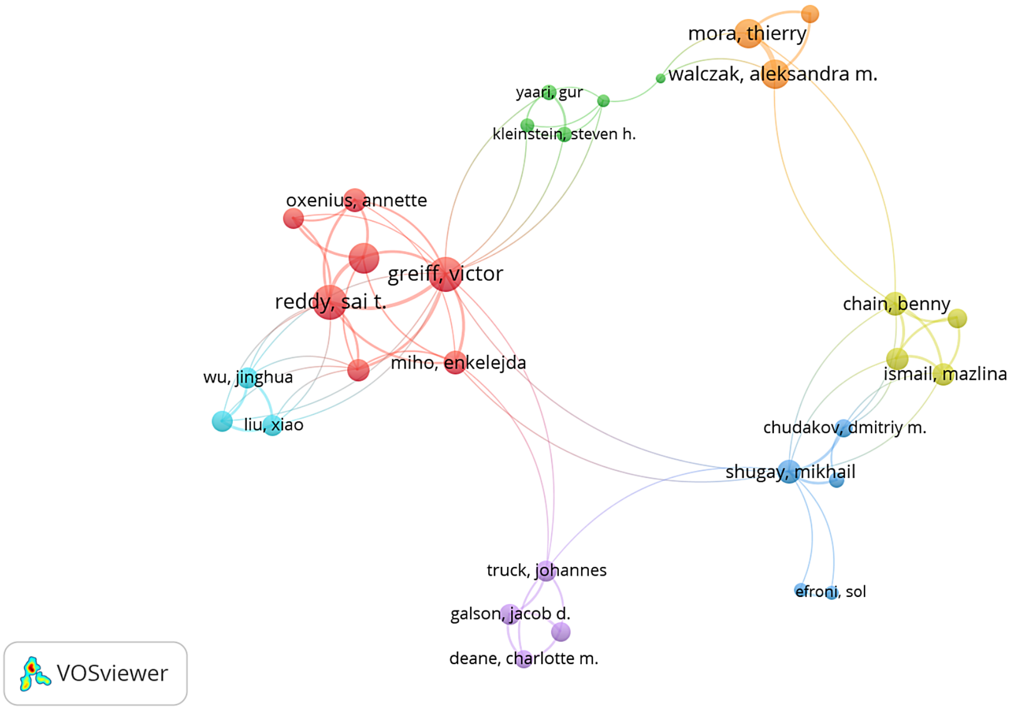

**Figure S5.** ***Co-authorship network of AIRR-simulation researchers.***
*This network was generated using the Total Link Strength analysis. The node size reflects the research output of each author, and the edge thickness indicates the collaboration strength. Color clusters identify major research communities: the red*

*cluster corresponds to computational immunology, the green cluster represents statistical physics approaches, and the connecting bridges highlight multidisciplinary collaborations.*

**Table S10.** Bibliometric profile of the top authors identified in the co-authorship network. *(Note: Authors are sorted by Total Link Strength)*

| Author | Documents | Citations | Total Link Strength |
|---|---|---|---|
| Greiff, Victor | 18 | 845 | 38 |
| Reddy, Sai T. | 14 | 657 | 38 |
| Yermanos, Alexander | 10 | 260 | 28 |
| Mora, Thierry | 19 | 656 | 26 |
| Walczak, Aleksandra M. | 19 | 656 | 26 |
| Chain, Benny | 11 | 308 | 18 |
| Shugay, Mikhail | 10 | 453 | 18 |
| Miho, Enkelejda | 7 | 390 | 17 |
| Oxenius, Annette | 5 | 58 | 17 |
| Ismail, Mazlina | 6 | 290 | 16 |
| Oakes, Theres | 6 | 288 | 16 |
| Weber, Cedric R. | 5 | 284 | 16 |

| |
|---|
| *(Truncated list for brevity; full data available in supplementary file)* |

#### S5.5 Co-citation Network Analysis

This section delineates the intellectual framework of the research on AIRR simulations. The co-citation network analysis, as illustrated in Figure S6, organizes frequently co-cited documents to uncover the field's conceptual underpinnings. Two predominant paradigms have been identified:

- Computational Immunology Paradigm (Red Cluster): High-throughput repertoire analysis and probabilistic modeling.
- Statistical Physics Paradigm (Green Cluster): Mathematical modeling and theoretical frameworks.

Victor Greiff appears as a central connector linking these two intellectual communities.

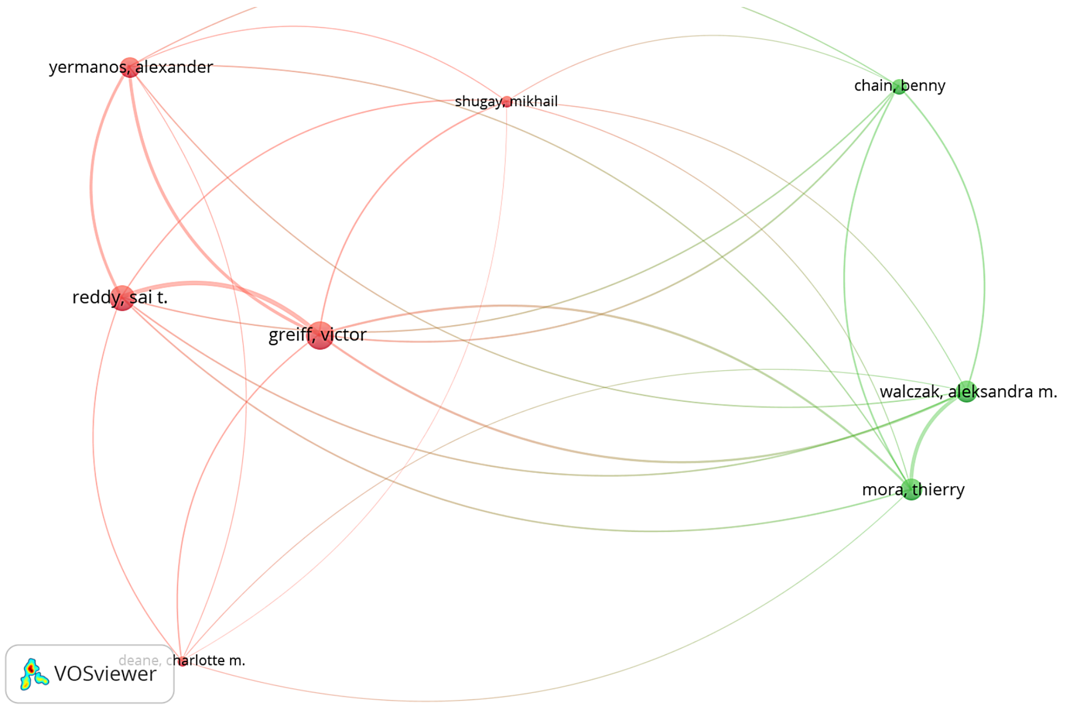

***Figure S6. Co-citation network analysis of key authors in AIRR-simulation research.***

*The node size reflects the total citation frequency, and the edge thickness represents the co-citation strength. Two major intellectual clusters emerge: (i) A computational-immunology cluster (red), centered around Alexander Yermanos, Sai T. Reddy, Victor Greiff, and Mikhail Shugay; and (ii) A statistical physics cluster (green), rooted in mathematical modeling. Greiff acts as a central connector between these paradigms.*

#### S5.6 Geographic and Institutional Collaboration Networks

The international collaboration landscape shows a strong geographic clustering.

- The United States operates as a major hub linking Europe and Asia.
- A robust transatlantic collaboration corridor connects the US with Switzerland, France, England, and Germany.
- Emerging research centers in Japan, China, and South Korea add to geographic diversity.

These findings are visualized in **Figure S7**.

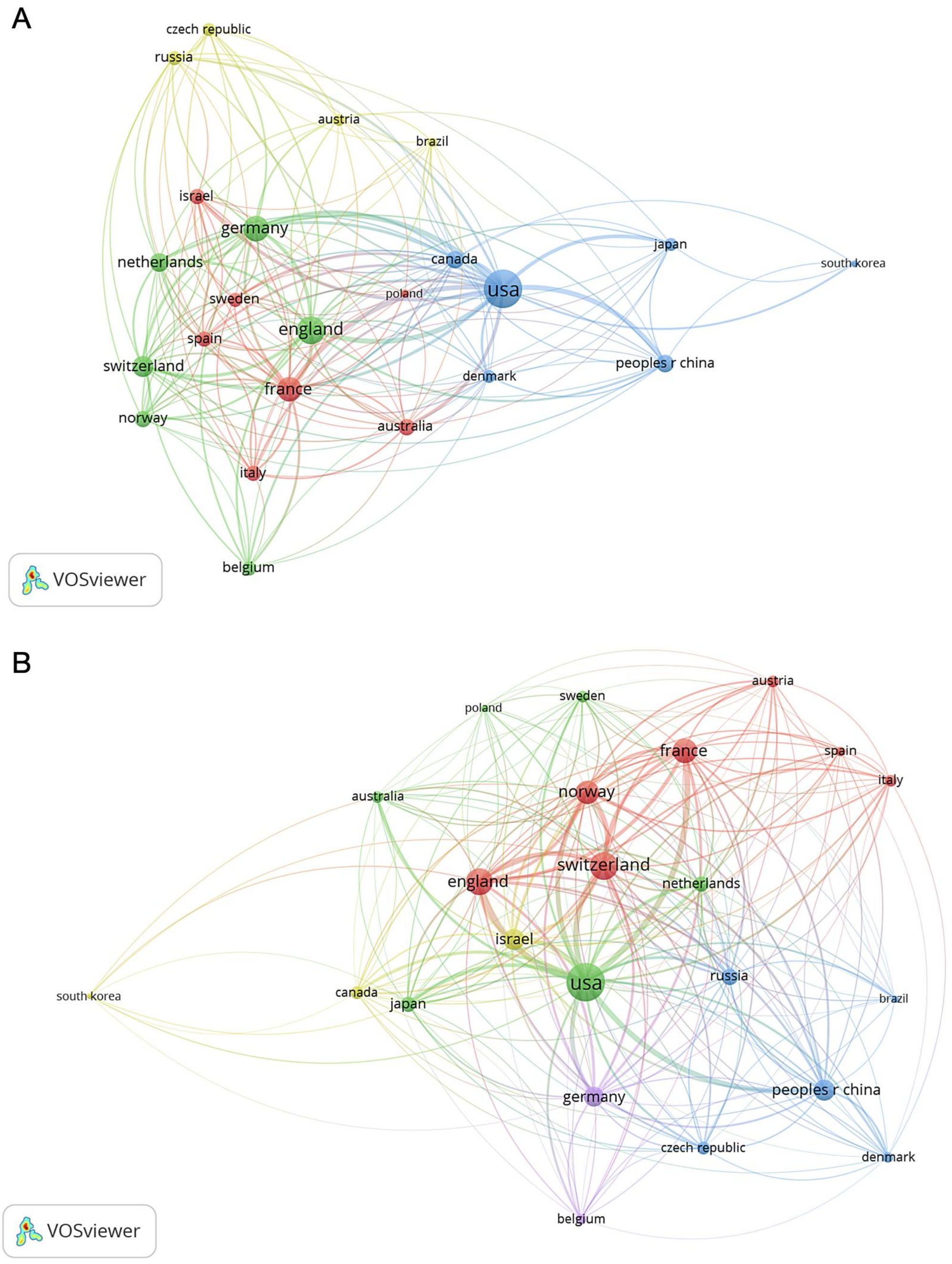

***Figure S7. Multi-dimensional analysis of global collaboration patterns in AIRR-simulation research (2015–2025).***

*(A) Country-level collaboration network: The USA is the dominant global connector. (B) Geographic co-citation heatmap: Strong intellectual proximity is observed between the USA, China, and major European countries, reflecting the convergence of methodological frameworks despite varying direct collaboration levels.*

#### S5.7 Strategic Implications and Future Directions

The integrated bibliometric analyses underscore several key priorities: (i) enhancing global coordination among both established and emerging research centers; (ii) promoting inter-paradigm collaboration between computational and theoretical frameworks; and (iii) building capacity through expanded training initiatives to reduce reliance on established hubs.

#### S5.8 Citation Trend Analysis

In addition to the network analyses, a citation trend analysis was conducted using Publish or Perish 8. The annual citation counts were aggregated to generate the trend curves shown in Figure S8. Notable findings include a consistent increase in citations from 2015 onwards and a marked acceleration in citations for machine learning-based tools (*e.g., TAPIR, TULIP*), alongside sustained citations for foundational frameworks.

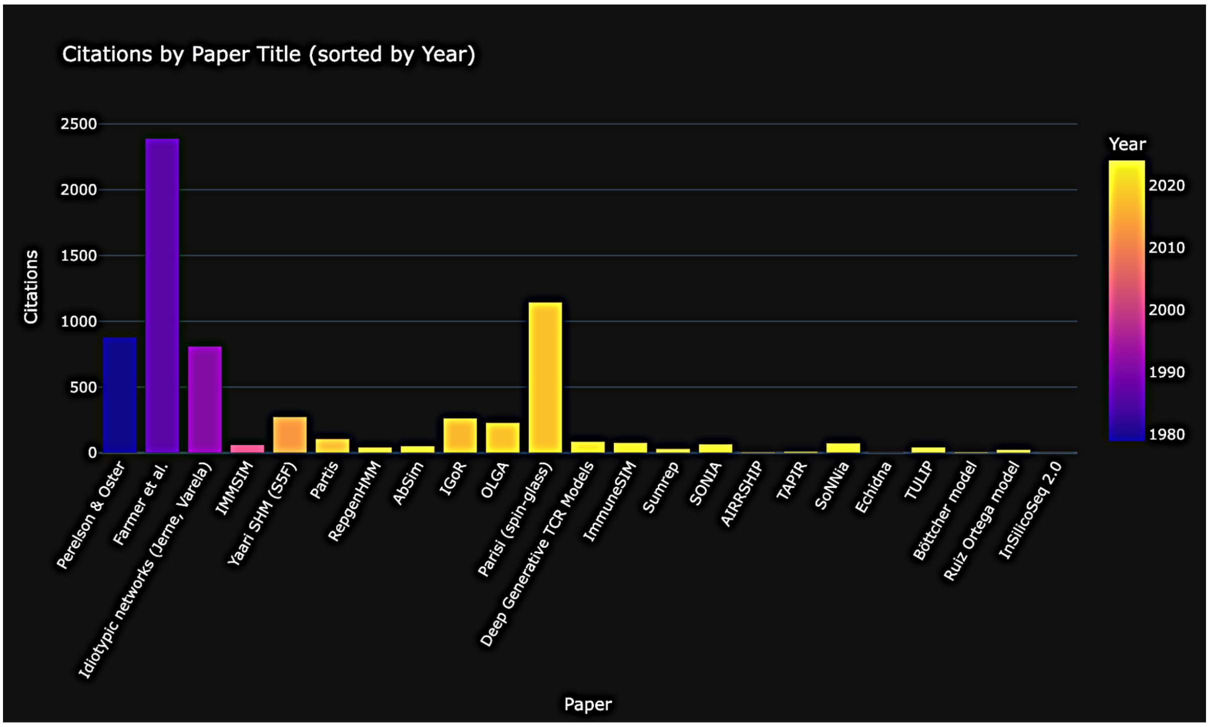

***Figure S8. Citation counts of major AIRR-simulation papers, sorted by publication year.***

*The chart highlights the historical shift from theoretical immune network models (blue–purple) to modern probabilistic and deep learning frameworks (yellow), which exhibit rapidly growing citation impact.*

***Table S12.*** *Structural limitations of current AIRR simulators and corresponding opportunities for improvement. This table summarizes the major conceptual, biological, and methodological gaps identified across widely used AIRR simulators. For each limitation, representative tools are listed alongside the resulting consequences for* *biological realism, inference accuracy, or reproducibility. The rightmost column outlines actionable directions for improving next-generation, context-aware, and modular AIRR-simulation frameworks.*

| Limitation | Representative Tools Affected | Consequences | Opportunities for Improvement |
|---|---|---|---|
| **Loss of biological context (spatial, temporal, functional)** | immuneSIM, AIRRSHIP, partis | Unrealistic lineage dynamics; inability to model tissue-specific responses | Incorporate spatial models, time-series data, and multi-omics integration |
| **Implicit, undocumented assumptions** | IGoR, OLGA, SONIA/soNNia | Hidden biases; inconsistent cross-tool interpretation | Explicit parameterization, open model specifications, provenance tracking |
| **Partial or missing ground truth** | immuneSIM (partial), AIRRSHIP | Limited lineage reconstruction benchmarking | Multi-scale ground truth: clonal labels, SHM paths, selection pressures |
| **Monolithic architectures** | Many classical simulators | Difficult to update or extend biological modules | Modular, declarative designs with replaceable components |
| **Limited interoperability with ML / multi-omics** | SONIA, IGoR, AbSim | Reduced applicability to translational modeling | Standardized APIs, AIRR-C compliance, ML-ready outputs |
| **Realism–scalability trade-off** | Detailed SHM/GC models vs. fast generators | Either slow execution or oversimplified biology | Hybrid multi-resolution models; GPU acceleration; cloud scaling |
| **Insufficient documentation or UI** | Several community tools | Barriers to adoption and reproducibility | Comprehensive docs, tutorials, containerization |

#### S5.7 AIRR Simulation Landscape Explorer

The AIRR Simulation Landscape Explorer is an interactive web-based tool developed to support navigation, comparison, and exploratory analysis of AIRR simulators within the UnivAIRRse framework. The explorer provides a two-dimensional visualization indexed by species (vertical axis) and publication year (horizontal axis), enabling users to dynamically filter simulators based on modeling approach, biological scope, abstraction level, chain support, output and ground-truth fidelity, use-case focus, and time-series capability.

The tool is publicly accessible via the IMGT® portal and can be used directly through a standard web browser without local installation. Hover-based interactive tooltips provide standardized metadata for each simulator, including modeled biological processes, supported receptor chains, output types, interface modality, species coverage, and intended applications. This functionality facilitates transparent comparison, reproducible tool selection, and rapid identification of simulators aligned with specific benchmarking or modeling objectives.

The AIRR Simulation Landscape Explorer is publicly available at:
https://www.imgt.org/AIRR-Simulator/
